# LOTUS: lineage-aware optimal transport reconstructs continuous clonal fate dynamics

**DOI:** 10.64898/2026.10.28.762359

**Authors:** Jingyuan Lu, Han Hou, Zhenyi Zhang, Yongjie Lin, Yihan Lin, Peijie Zhou

## Abstract

Destructive sampling in single-cell sequencing obscures ancestor–descendant relationships, limiting the continuous reconstruction of cell-fate dynamics. Here we present Lineage-aware optimal transport for unbalanced single-cell dynamics (LOTUS), a framework that integrates time-resolved transcriptomic snapshots with clonal lineage information from engineered barcodes or endogenous T cell receptor clonotypes. LOTUS couples clone-level optimal transport with neural unbalanced transport to model continuous cellular state change, population growth and clonal propagation under lineage-consistent constraints. In branching simulations, LOTUS reduced lineage-inconsistent transport relative to benchmark methods, lowering off-target routing from 4.94–14.84% to 1.10%. On LARRY lineage-barcode scRNA-seq of hematopoiesis, LOTUS reconstructed branching trajectories, improved early neutrophil–monocyte fate-boundary prediction, and inferred multi-fate progenitors and fate-biased meta-clones, linking their distinct fate potentials to inferred epigenetic priming. On CellTagging scRNA-seq of fibroblast reprogramming, it distinguished reprogrammed from failed trajectories and prioritized fate-associated perturbation genes. In longitudinal scRNA-seq and TCR-seq of malaria-responsive memory (CD45RA-) CD4+ T cells, it preserved endogenous clonotype structure and reconstructed TR1 dynamics and regulatory networks. LOTUS thus converts lineage-resolved single-cell snapshots into interpretable, continuous models of fate dynamics and their regulatory programs.

## Introduction

Cell-fate decisions underlie the dynamic behavior of multicellular systems, including embryonic development, tissue homeostasis, injury repair, cellular reprogramming and immune responses^1–3^. During these processes, cells continuously change their transcriptional state, expand clonally and progressively restrict their fate potential, ultimately giving rise to specialized cell types. Understanding how early cellular states are linked to later fate outcomes is therefore central to developmental regulation, regeneration and disease progression^2,4^.

Single-cell RNA sequencing (scRNA-seq) measures gene expression across large numbers of individual cells^5^ and, together with trajectory-inference methods, enables the reconstruction of continuous state-transition processes from discretely sampled populations^6–8^. Because sequencing destroys the profiled cells, however, scRNA-seq cannot directly track the same cell from an early state to its later fate^9,10^. Trajectory-inference methods therefore typically rely on transcriptomic similarity, local neighborhood structure or distribution matching to infer dynamic relationships among cells. Yet transcriptomic similarity does not necessarily reflect lineage history^11^: cells can retain fate-relevant molecular memory along lineages that is not captured by the instantaneous transcriptome, including heritable or otherwise stable features^12–14^. Such memory can bias fate potential, allowing early progenitors that appear similar in expression space to differentiate toward distinct terminal fates. Consequently, dynamical reconstruction based solely on transcriptomic geometry can yield state transitions that are inconsistent with clonal propagation, compromising the inference of fate boundaries, branching structure and regulatory mechanisms.

Lineage-tracing single-cell sequencing (LT-scSeq) provides a key additional layer of information with which to address this problem^1,2,4,15,16^. By measuring clonal identity together with transcriptional state over time, LT-scSeq links cellular state to clonal history at single-cell resolution. Lineage information can be obtained from exogenous barcoding systems that either label progenitors once or record accumulated editing events, as well as from endogenous markers such as shared immune-receptor clonotypes^17–21^. Here we focus on time-resolved single-cell datasets in which clonal identity is encoded by stable lineage labels, including experimental barcodes from systems such as LARRY^11^, CellTagging^22^ and Polylox^23^, and endogenous clonotypes. As cells proliferate and differentiate, these labels are transmitted to descendants, so that cells sharing a common ancestor can be identified at later time points and clonal fate bias and differentiation output can be quantified **(Fig. 1a).** Because such datasets jointly measure transcriptional state, sampling time and clonal identity, they provide clone-level constraints for reconstructing lineage-consistent cell-state transitions **(Fig. 1b).**

**Fig. 1.**
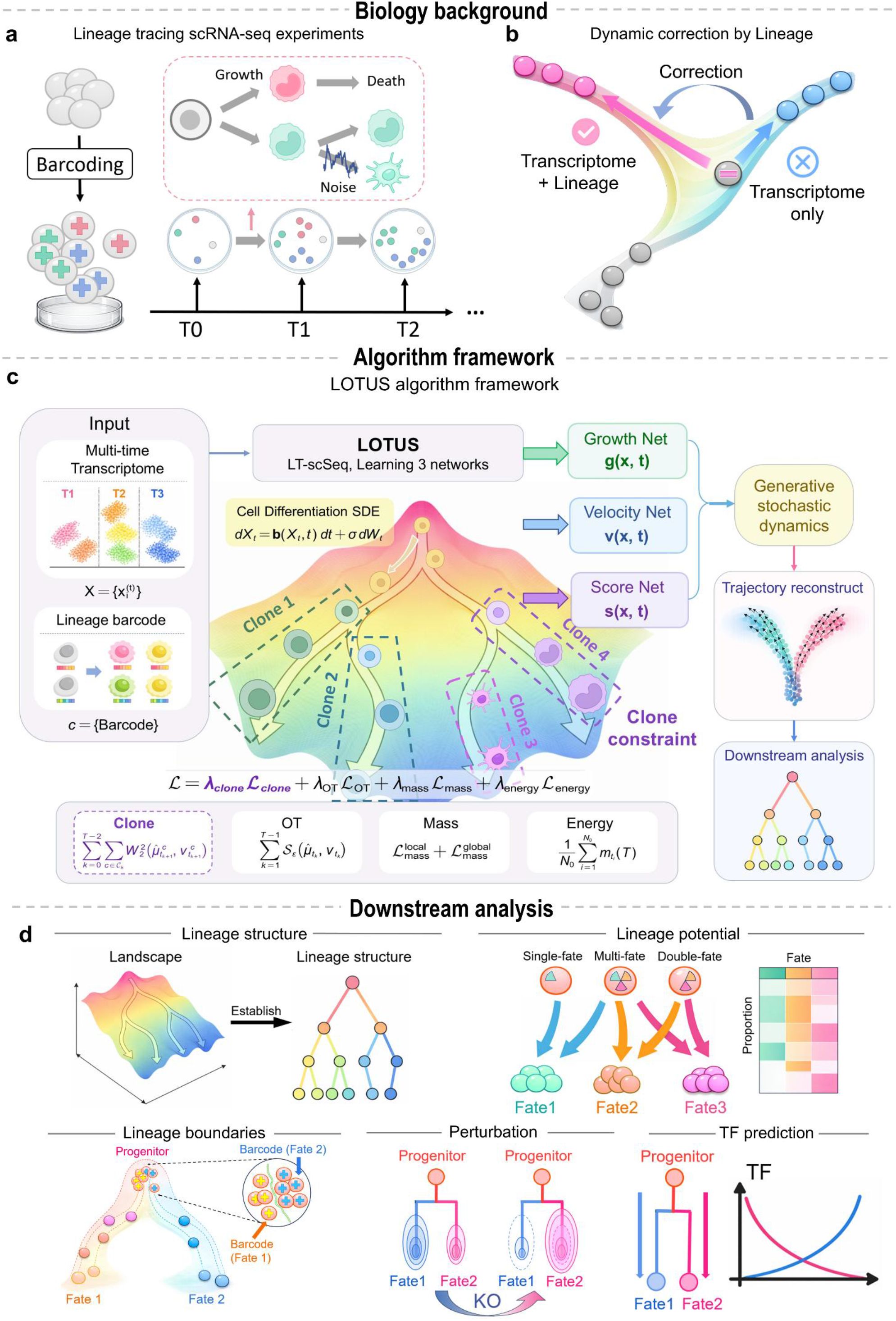
Overview of the LOTUS framework and its applications. **a,** Schematic illustration of lineage tracing single-cell sequencing (LT-scSeq) experiments. Cells are barcoded at the initial stage and profiled across multiple time points. **b,** Lineage information provides an additional constraint for correcting transcriptome-inferred dynamics. Compared with transcriptome-only reconstruction, integrating lineage barcodes helps distinguish transcriptionally similar but lineage-inconsistent transitions and improves the recovery of biologically plausible developmental trajectories. **c,** Algorithmic framework of LOTUS. LOTUS integrates time-resolved single-cell transcriptomes with clonal barcode information to infer lineage-constrained cell-state dynamics. The model formulates population evolution as a lineage-aware regularized unbalanced optimal transport problem and is optimized within a neural probability-flow framework. Three neural networks parameterize the probability-flow velocity v(x,t), effective net growth rate g(x,t) and scaled log-density s(x,t). Training combines distributional optimal transport matching, mass matching, energy regularization and clone-level optimal transport, with clonal constraints applied to probability-flow ODE predictions. The learned fields also support stochastic trajectory generation. **d,** Downstream analysis tasks. LOTUS can be used to infer cell-fate landscapes, establish lineage structures, estimate fate potentials, identify lineage boundaries, simulate perturbation effects and predict transcription factors associated with fate transitions.

However, lineage-resolved single-cell data are typically sparse, heterogeneous and incompletely sampled: lineage labels are detected only in a subset of cells, clone sizes vary by orders of magnitude, and each time point captures only part of the underlying population^24^. The central computational challenge is therefore to infer continuous evolution through transcriptional space while using clonal history to constrain cross-time state changes.

Several methods have integrated clonal lineage information with single-cell states to infer cell fate^4,25^. CoSpar^24^ reconstructs transition maps from sparse clonal observations, whereas Super-OT^26^, LineageOT^27^ and moslin^28^ extend optimal transport frameworks to incorporate lineage information, yielding probabilistic ancestor–descendant couplings. DestinyNet^29^ learns a fate-aware latent space from clonal triplets to predict early fate bias. CLADES^30^ estimates clone-level proliferation and differentiation rates, whereas CORAL^31^ identifies heritable transcriptional states. Despite their complementary strengths, these lineage-aware methods generally represent fate dynamics as transition maps, coupling matrices, lineage trees, clone-level rate parameters or discrete heritable states. These methods do not formulate cell-state evolution as a continuous-time generative process directly constrained by observed clonal propagation.

Conversely, neural dynamical models and unbalanced optimal transport methods, including TIGON^32^, DeepRUOT^33,34^ and scDiffEq^35^, can reconstruct continuous trajectories from time-series single-cell data and model non-conserved cellular mass arising from proliferation, cell loss or sampling variation. These models, however, are constrained only by global distribution matching or transcriptomic geometry and do not use observed clonal propagation to regularize cross-time state transitions. They can therefore produce trajectories that are plausible in transcriptional space but inconsistent with clone-level lineage relationships. Reconstructing lineage-resolved cell-fate dynamics thus requires a model that jointly learns continuous transcriptional state evolution, non-conserved cellular mass and clonal consistency over time.

Here we present Lineage-aware optimal transport for unbalanced single-cell dynamics (LOTUS), a framework for reconstructing continuous cell-fate dynamics from time-resolved, lineage-tracing single-cell data. LOTUS formulates cell-population evolution as a lineage-regularized unbalanced optimal transport problem within a neural probability-flow framework. By coupling neural unbalanced transport with a clone-level constraint on observed clonal propagation, LOTUS jointly learns continuous transcriptional state dynamics, non-conserved cellular mass and lineage-consistent clone propagation. This unified framework enables lineage-aware inference of early fate bias, differentiation boundaries and clone-associated regulatory programs, and supports in silico perturbation-based prioritization of candidate regulators.

## Results

### LOTUS embeds clonal lineage constraints in neural optimal transport dynamics

To reconstruct lineage-resolved cell-fate dynamics from destructive single-cell measurements, we developed Lineage-aware optimal transport for unbalanced single-cell dynamics (LOTUS), a framework that integrates time-resolved single-cell transcriptomes with clonal barcode information **(Fig. 1a and Methods).** LOTUS combines two complementary sources of information: global transcriptomic distributions, which describe population-level changes in cell state, and shared clonal barcodes, which constrain the inferred transport dynamics to remain consistent with observed clonal propagation.

LOTUS represents an evolving cell population as a time-dependent density in a PCA-reduced transcriptomic space. To model cell-state transitions and changes in cell abundance jointly, we formulated the problem as a regularized unbalanced optimal transport problem with stochasticity and growth. In this formulation, deterministic cell-state changes are parameterized by a neural velocity field, and local expansion or depletion is captured by a neural growth field. Starting from cells sampled at an observed time point, LOTUS propagates cell states to later time points through a neural ODE with inferred mass changes. LOTUS can thus generate continuous cell-state trajectories, estimate local mass expansion or depletion and interpolate unobserved intermediate states between measured time points.

The LOTUS objective combines four complementary losses **(Fig. 1c).** A clone-level optimal transport loss provides the lineage constraint by promoting transport between barcode-sharing cells across time points. A Sinkhorn reconstruction loss aligns the predicted and observed target distributions, a mass-matching loss links the learned growth field to changes in cell abundance, and an energy-regularization term imposes a minimum-action constraint that encourages smooth, low-energy trajectories. For each pair of adjacent time points, LOTUS identifies barcodes shared between the source and target populations and computes, within each shared clone, an optimal transport distance between predicted descendants and observed target cells carrying the same barcode. This loss penalizes dynamics that match the global distribution but route clonal descendants into transcriptomic regions inconsistent with their observed lineage output, a failure mode that is particularly likely when distinct lineages pass through nearby or overlapping regions of transcriptomic space: without lineage information, multiple transport plans may appear equally plausible near branching regions or early progenitor states despite implying different clonal histories **(Fig. 1b).** After training, LOTUS yields continuous lineage-constrained trajectories together with local mass-change estimates and clone-resolved transport patterns, supporting downstream analyses such as fate-boundary inference, clone-level fate-potential estimation and in silico perturbation-based regulator prioritization **(Fig. 1d).**

### Benchmarking on simulated datasets

We first evaluated whether LOTUS preserves lineage-consistent fate assignment using simulated lineage-tracing data with known barcode–fate relationships. For simulated dataset 1, we constructed a two-time-point developmental model with a bifurcating fate structure: at the first time point, two progenitor populations occupied nearby regions of transcriptomic space but carried distinct barcodes; by the second time point, they had differentiated into two distinct terminal fates. This design tests whether a model can correctly route cells to the barcode-associated fate branch even when early progenitor states are transcriptionally similar. The off-target rate captured two types of reconstruction error: cross-branch misrouting, in which predicted cells entered the wrong terminal branch, and neighborhood deviation, in which predicted cells remained far from the terminal fate associated with their barcode.

Across ten independently generated test datasets, LOTUS achieved the lowest mean off-target rate (1.10%), compared with DeepRUOT (5.20%), TIGON (4.94%) and scDiffEq (14.84%) **(Fig. 2b).** Representative reconstructions from one test dataset further showed that LOTUS more faithfully followed the barcode-defined fate branches, whereas the baseline methods exhibited varying degrees of cross-branch transport and displacement from the observed terminal states **(Fig. 2c–f).** LOTUS also yielded the lowest mean final-time-point Wasserstein-2 distance between predicted and observed distributions (1.41 versus 1.76 for DeepRUOT, 1.66 for TIGON and 2.33 for scDiffEq; **Supplementary Fig. 1e**). Together, these results show that incorporating clone-level constraints improves both lineage consistency and terminal-state reconstruction.

**Fig. 2.**
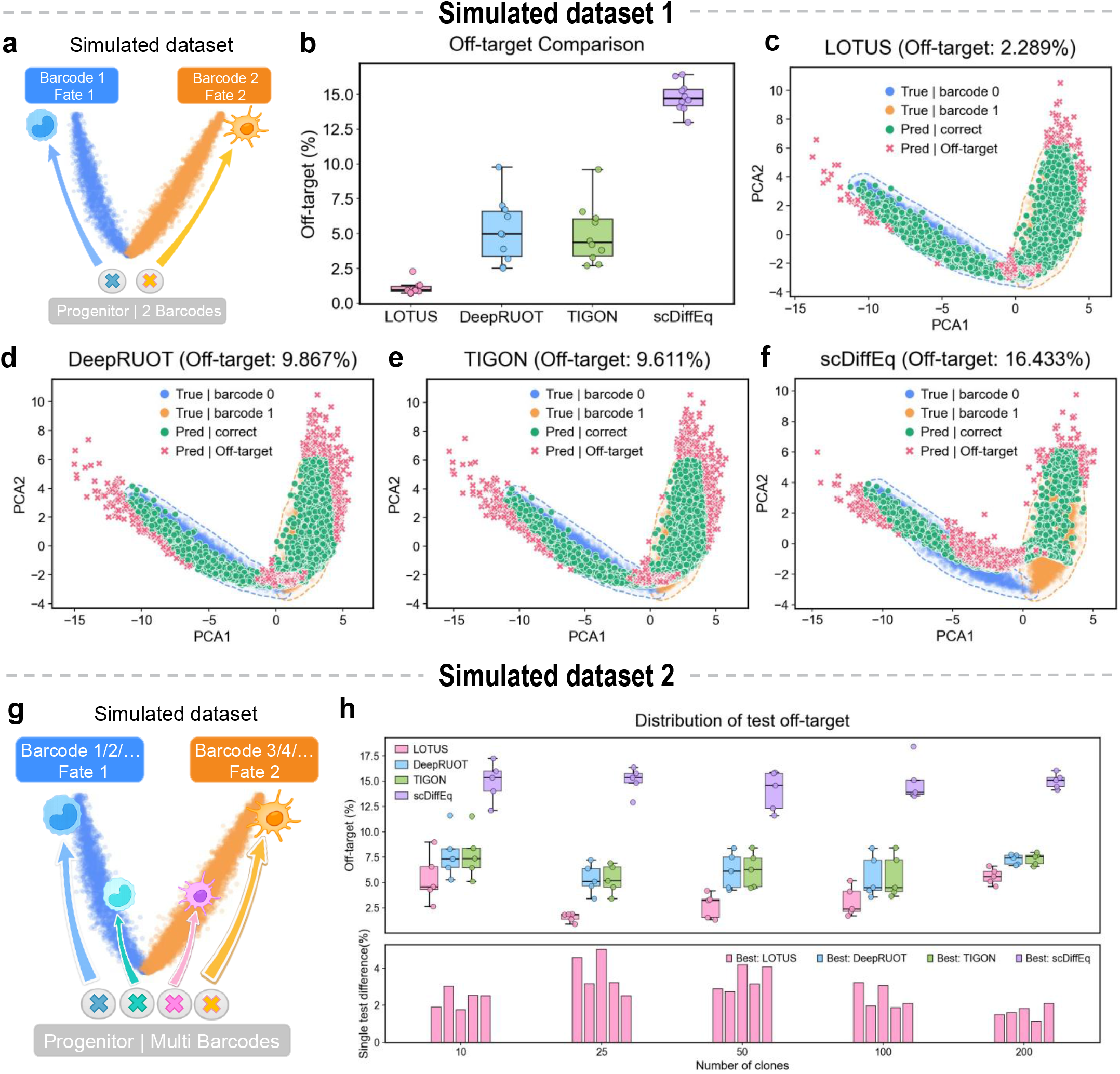
LOTUS improves lineage-consistent trajectory reconstruction in simulated lineage-tracing datasets. **a,** Schematic of simulated dataset 1, in which two barcoded progenitor populations give rise to two distinct terminal fates. This setting was designed to evaluate whether each method can preserve barcode-specific fate assignment during trajectory reconstruction. **b,** Quantification of off-target rates across repeated simulations for LOTUS, DeepRUOT, TIGON and scDiffEq. Each point represents an independent test run performed on a newly generated simulated dataset. The off-target rate represents the proportion of generated cells that fall outside the neighborhood of their corresponding true terminal branch. LOTUS shows a lower fraction of off-target cells than the baseline methods. Boxes indicate the interquartile range, center lines indicate the median and points represent individual simulation runs. **c–f,** PCA visualizations of reconstructed trajectories for LOTUS (**c**), DeepRUOT (**d**), TIGON (**e**) and scDiffEq (f) in one representative test dataset of simulated dataset 1; annotated values refer to this single dataset. True cells from the two barcode-associated fates are shown in blue and orange, correctly routed predicted cells are shown in green, and off-target predictions are shown in red. Dashed contours indicate the expected fate regions. **g,** Schematic of simulated dataset 2, in which multiple barcoded progenitor populations give rise to two distinct terminal fates. Compared with simulated dataset 1, this dataset contains more clones. This setting evaluates the robustness of lineage-aware reconstruction under increasing clonal complexity. **h,** Distribution of test off-target rates across different numbers of clones. The upper panel shows the distribution of off-target rates for LOTUS, DeepRUOT, TIGON and scDiffEq under each clone number, where each point represents an independent test run performed on a newly generated simulated dataset. The lower panel shows, for each test dataset, the performance margin between the top-ranked and second-ranked methods, computed as the second-lowest off-target rate minus the lowest off-target rate. LOTUS maintains lower off-target rates across clone numbers compared with DeepRUOT, TIGON and scDiffEq.

We next tested the robustness of LOTUS in a more clonally complex setting, in which multiple progenitor clones carrying distinct barcodes generated descendants in parallel along the same bifurcating process, giving rise to two terminal fates. This setting tests the ability of each model to preserve lineage-consistent transport as clonal complexity increases **(Fig. 2g).** As the number of clones increased from 10 to 200, LOTUS maintained a low off-target rate and consistently outperformed DeepRUOT, TIGON and scDiffEq **(Fig. 2h, top, and Supplementary Fig. 1a-d).** To quantify this advantage, we calculated the margin between the lowest and second-lowest off-target rates for each setting: LOTUS was consistently the top-performing method and typically maintained a positive margin over the next-best method **(Fig. 2h, bottom).** Together, these simulations show that LOTUS preserves barcode-associated fate assignment in bifurcating systems and remains robust as clonal complexity increases, reducing lineage-inconsistent transport in multi-clone differentiation settings.

### LOTUS reconstructs lineage-resolved hematopoietic trajectories and early fate boundaries

To assess whether LOTUS can recover continuous differentiation trajectories from multi-time-point LT-scSeq data, we applied it to the LARRY hematopoietic differentiation dataset^11^, which jointly profiles single-cell transcriptomes and clonal lineage barcodes during in vitro hematopoietic differentiation, spanning the progression from undifferentiated progenitor-like states toward multiple hematopoietic lineages. In the SPRING embedding, cells formed a branched differentiation manifold, with undifferentiated progenitor-like cells located centrally and branches extending toward neutrophil, monocyte and other hematopoietic states **(Fig. 3b).**

**Fig. 3.**
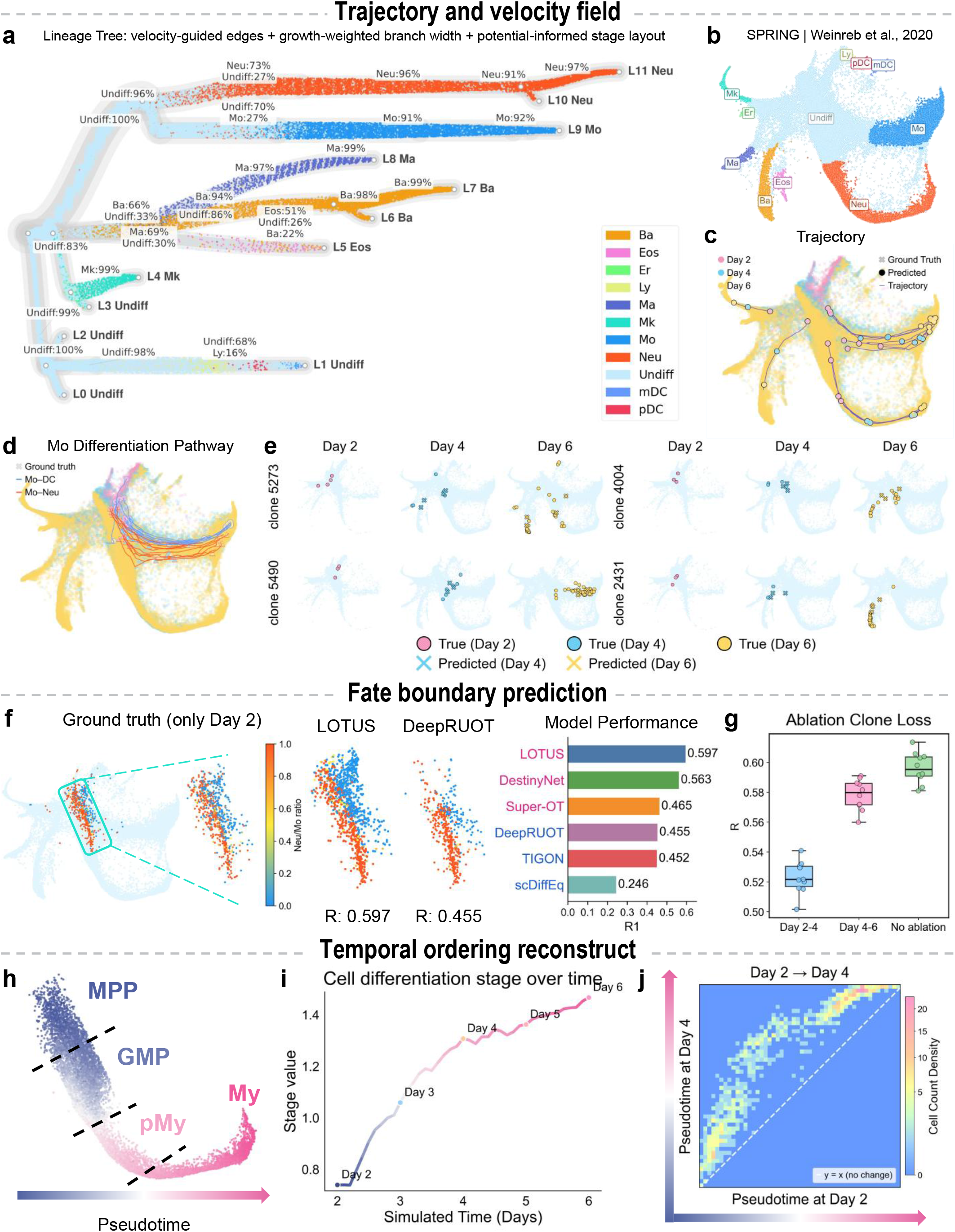
Temporal order reconstruction of lineage-resolved differentiation trajectories and fate-boundary dynamics. **a,** LOTUS-derived lineage-tree representation of LARRY hematopoietic differentiation. Barcoded cells were grouped into PCA-based Leiden microclusters and arranged along a LOTUS potential–derived developmental stage, with the undifferentiated-enriched microcluster used as the root. Candidate microcluster connections were obtained from the PCA-neighbor graph, constrained to proceed from earlier to later stages, and refined by LOTUS-inferred velocity alignment to favor forward developmental transitions. Cells were projected onto the resulting root-to-leaf paths and colored by annotated cell state. Branch widths encode downstream growth-weighted branch strength based on subtree cell abundance and LOTUS-inferred growth rate. Lineage-end labels indicate terminal dominant states, and local labels show dominant cell-state compositions within lineage-stage bins. **b,** SPRING visualization of the LARRY scRNA-seq hematopoietic differentiation dataset, colored by cell type annotations. **c,** LOTUS-inferred trajectory reconstruction on the SPRING embedding. Ground-truth cells at different time points are shown together with model-predicted cell states and sampled stochastic trajectories, illustrating the ability of the model to recover continuous developmental paths across discrete sampling days. **d,** Reconstruction of monocyte-associated differentiation routes. Inferred trajectories capture divergent paths from early progenitor states toward Mo–DC and Mo–Neu branches, consistent with the topology of the reference manifold. **e,** Representative clone-level dynamics across day 2, day 4 and day 6. True clone distributions and predicted future states are shown for individual barcoded clones, demonstrating clone-specific propagation along lineage-consistent differentiation paths. **f,** Fate-boundary prediction between neutrophil and monocyte fates. Ground-truth fate ratios for Day 2 cells and predictions by LOTUS and DeepRUOT are shown for the highlighted region. Colors indicate the neutrophil-to-monocyte fate ratio. Bar plots show Pearson correlation *R* with the ground truth for LOTUS, DestinyNet, Super-OT, TIGON, DeepRUOT and scDiffEq. **g,** Ablation of temporal lineage supervision. Pearson correlation with the ground-truth fate boundary after removing the Day 2–4 or Day 4–6 clone-loss constraint. Early lineage supervision had a greater contribution to fate-boundary prediction. **h,** Schematic representation of pseudotime progression along the myeloid differentiation continuum, spanning MPP, GMP, pMy and My states. **i,** Reconstructed temporal ordering of cell differentiation stages over simulated time. For each simulated time point, the stage value was calculated as the average ordinal differentiation-stage score of cells, with MPP, GMP, pMy and My encoded as stages 0-3, respectively. The inferred stage value increases from early to late simulated days, indicating that the learned dynamics are temporally consistent with progressive myeloid differentiation. **j,** Pseudotime progression from day 2 to day 4. Each cell was assigned a pseudotime value at the observed day 2 state and predicted day 4 state by weighted nearest-neighbor mapping in PCA space to reference cells with known pseudotime. The two-dimensional density map shows the paired pseudotime values of the same cells, with the x axis indicating pseudotime at day 2 and the y axis indicating pseudotime at day 4. The dashed diagonal denotes no pseudotime change. Enrichment of density above the diagonal indicates that most cells acquire higher pseudotime values from day 2 to day 4, consistent with forward progression along the inferred developmental trajectory.

To summarize the global dynamics learned by LOTUS, we integrated the inferred velocity, growth and potential information into a lineage-tree representation. The tree originated from the undifferentiated-cell-enriched region and extended toward multiple terminal branches along the inferred developmental progression; branch orientation was supported by the LOTUS velocity field, and branch width reflected growth-weighted downstream branch strength. This representation recovered the hierarchical organization of hematopoietic differentiation, including trajectories from undifferentiated progenitor states toward neutrophil, monocyte, basophil, mast-cell and megakaryocyte-associated branches, consistent with the established hematopoietic hierarchy^36^ **(Fig. 3a).** The dominant cell-state composition of terminal branches agreed with their annotated fates, indicating that LOTUS not only connects cell states across discrete time points but also organizes continuous dynamics into a directional, branch-resolved lineage structure.

At the trajectory level, LOTUS connected discrete sampling time points into continuous developmental paths across the SPRING manifold. Predicted states and sampled stochastic trajectories progressed from the day 2 early progenitor region through intermediate day 4 states toward multiple day 6 terminal branches **(Fig. 3c).** This trajectory structure is consistent with the observed temporal ordering and branching topology, suggesting that the learned velocity field captures a global developmental flow from early progenitors to terminal hematopoietic lineages. Denser trajectory sampling and velocity streamlines further supported this interpretation, with streamlines extending from the high-potential early region toward lower-potential differentiation branches and separating at major branch points **(Supplementary Fig. 2a–c).** Consistent with the two monocyte differentiation routes previously linked to distinct clonal coupling with DC or Neu fates in LARRY^11^, LOTUS reconstructed spatially distinct trajectories toward Mo–DC and Mo–Neu directions (Fig. 3d), indicating that the model resolves lineage-origin differences within a single annotated cell type. In parallel, the spatial pattern of LOTUS-inferred growth was concordant with proliferation-related cell-cycle signals, suggesting that the model captures state transitions together with local mass-change patterns **(Supplementary Fig. 2d,e).** At the clone level, predicted day 4 and day 6 descendants of representative barcoded clones followed the same distributional directions as their observed clone members while preserving clone-specific expansion toward different fate regions **(Fig. 3e).**

We next examined whether LOTUS could recover early fate bias at the neutrophil–monocyte boundary **(Fig. 3f, Supplementary Fig. 3a).** The LARRY dataset was partitioned into clone-disjoint training and test subsets with similar transcriptomic distributions, and day 2 differentiation bias was defined by mapping the proportions of day 4/6 neutrophil and monocyte cells back to their corresponding clones. In held-out test cells, LOTUS reconstructed a fate-ratio distribution that closely matched the reference boundary: Neu-biased and Mo-biased cells were separated along the early differentiation region while retaining a continuous transition between the two fate biases. Quantitatively, LOTUS achieved the highest agreement with the reference fate ratio (*R* = 0.597), followed by the lineage-informed methods DestinyNet (*R* = 0.563) and Super-OT (*R* = 0.465), and outperformed TIGON (*R* = 0.452), DeepRUOT (*R* = 0.455) and scDiffEq (*R* = 0.246), suggesting that incorporating lineage information generally improves early fate-boundary reconstruction. Notably, this improvement was preserved for clones whose lineage information was entirely withheld from training, indicating that clone supervision generalizes to unseen clones rather than memorizing clone outputs. To include CoSpar as a lineage-aware transition-map baseline, we repeated the Neu–Mo fate-boundary analysis using all available cells. In this full-dataset setting, LOTUS again showed the strongest agreement with the reference boundary (*R* = 0.586), exceeding DeepRUOT (*R* = 0.487) and CoSpar (*R* = 0.431; **Supplementary Fig. 3b**). In a global cell-level fate-direction prediction task, LOTUS also better recovered reference differentiation directions derived from Weinreb et al. than DeepRUOT (accuracy = 0.720 versus 0.675; Supplementary Fig. 4).

To determine how lineage constraints from different temporal intervals contribute to fate-boundary inference, we performed temporal ablation experiments **(Fig. 3g).** Removing either the day 2–4 or the day 4–6 clone loss preserved the broad spatial trend of the Neu/Mo fate boundary but reduced agreement with the reference boundary to mean *R* = 0.523 and mean *R* = 0.578, respectively. The larger decrease after removing the day 2–4 constraint indicates that early clonal information is particularly important for resolving fate bias, consistent with the notion that hematopoietic progenitors can carry future fate tendencies before terminal phenotypes fully separate.

Finally, we evaluated whether LOTUS preserves the temporal ordering of hematopoietic differentiation. Following the neutrophil differentiation pseudotime axis, we summarized myeloid maturation as four consecutive stages, MPP, GMP, pMy and My, encoded as stage values 0–3, respectively **(Fig. 3h).** Along LOTUS-simulated time, the average stage value increased continuously from day 2 to day 6, with a gradual reduction in slope at later times, consistent with progressive maturation from early progenitor states toward more differentiated myeloid states **(Fig. 3i).** At the single-cell level, paired pseudotime values for observed day 2 cells and their predicted day 4 states were concentrated above the diagonal, indicating that most cells advanced along pseudotime during forward prediction **(Fig. 3j).** Together, these results show that LOTUS reconstructs not only branching fate structure and fate-boundary organization but also the directional temporal progression of hematopoietic differentiation.

### LOTUS supports lineage-aware in silico perturbation analysis in LARRY hematopoiesis

Generative models of cellular state dynamics have been used for in silico perturbation analysis, in which transcriptional changes in early progenitors are used to estimate shifts in downstream state transitions and terminal fate composition^37,38^. Motivated by these studies, we asked whether the lineage-aware dynamics learned by LOTUS could support in silico perturbation analysis of how gene-level changes in early progenitors shift predicted downstream fate allocation. We first focused on the LARRY dataset, in which barcoded day 2 progenitors give rise to multiple myeloid terminal fates by day 6. As a proof of concept, we perturbed day 2 progenitors along PCA-loading directions associated with *Cebpe, Mxd1* and *Dach1*, genes linked to granulocytic maturation and myeloid differentiation programs, and simulated their subsequent differentiation trajectories using the learned LOTUS dynamics^39,40^. Compared with the unperturbed control, perturbed trajectories showed a stronger bias toward the neutrophil branch, with an increased predicted day 6 neutrophil fraction and a concomitant decrease in predicted monocyte fraction **(Fig. 4a and Supplementary Fig. 5a,b).** A perturbation-strength analysis further showed that increasing positive perturbation strength led to a dose-dependent increase in predicted neutrophil fraction and a corresponding decrease in monocyte fraction, whereas negative perturbations induced the opposite fate shift, indicating that the perturbation responses captured by LOTUS are continuous and directionally consistent **(Supplementary Fig. 5c).**

**Fig. 4.**
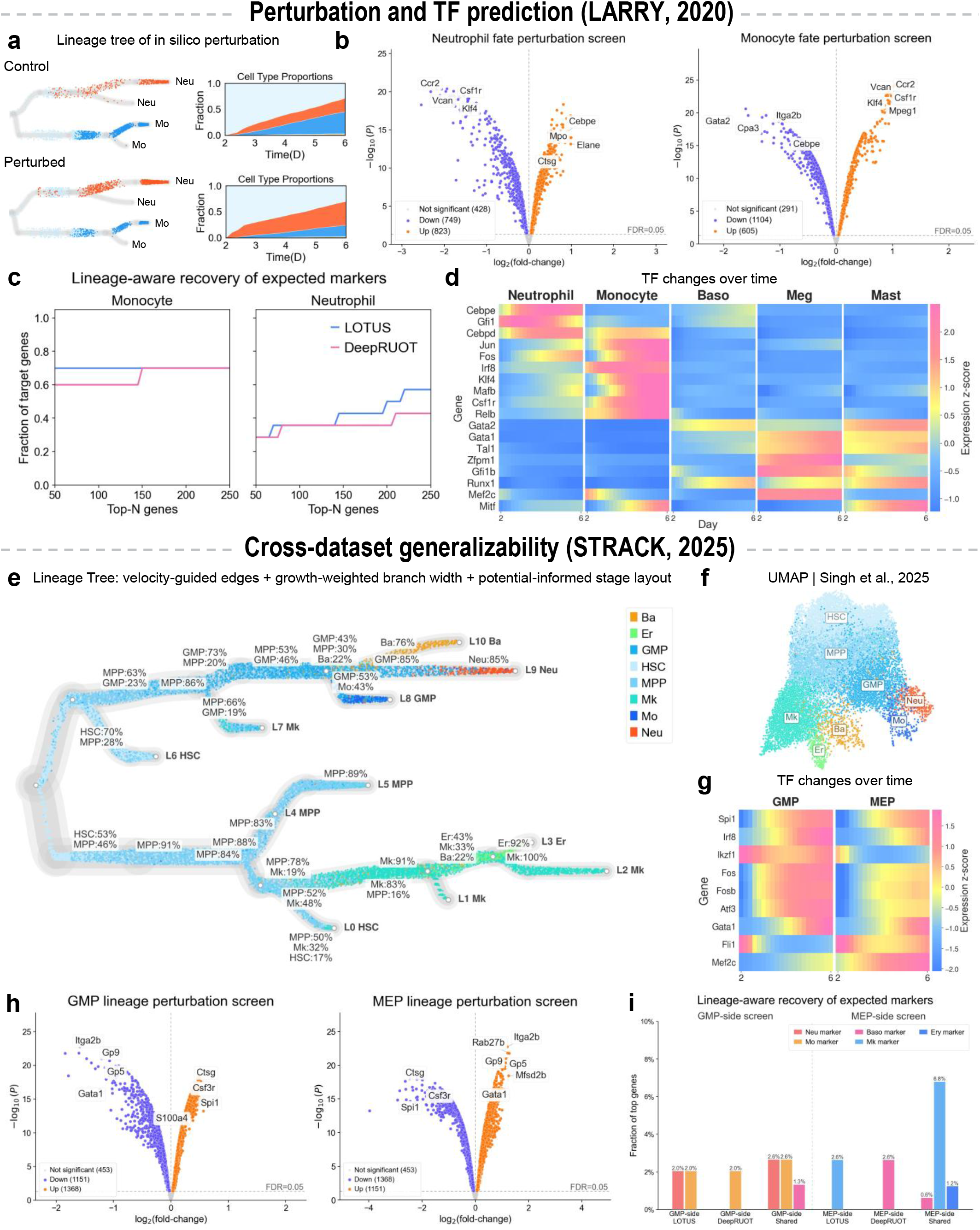
Lineage-aware *in silico* perturbation analysis and transcription-factor prediction across datasets. **a–d** show *in silico* perturbation analyses and transcription-factor dynamics based on the LARRY hematopoietic differentiation dataset from Weinreb et al. (2020). **a,** *In silico* perturbation of neutrophil–monocyte fate allocation in the LARRY dataset. The perturbation was applied to Cebpe, Mxd1 and Dach1: in the perturbation input, day-2 progenitor cells were displaced in PCA space along the PCA-loading directions of these genes, while the control input was generated without gene perturbation. Lineage trees show control and perturbed trajectories, with neutrophil and monocyte branches highlighted in red and blue, respectively. Stacked area plots quantify the corresponding temporal changes in cell-type proportions from day 2 to day 6. **b**, Volcano plots showing gene perturbation effects on neutrophil and monocyte fate probabilities. Genes with significant positive and negative effects are highlighted in orange and purple, respectively, whereas non-significant genes are shown in grey. Representative lineage-associated regulators are annotated. The dashed horizontal line denotes the FDR threshold. **c**, Fraction of predefined lineage markers and transcription factors recovered among the top-ranked perturbation genes for monocyte- and neutrophil-associated screens. Lines show recovery across Top-N genes (N = 50–250) for LOTUS and DeepRUOT. **d**, Temporal expression dynamics of transcription factors associated with major hematopoietic lineages in the LARRY dataset. Heatmap values represent gene-wise expression z-scores across inferred differentiation time, grouped by neutrophil, monocyte, basophil, megakaryocyte and mast-cell programs. **e–i** show lineage-tree reconstruction, *in silico* perturbation analyses and transcription-factor dynamics based on the STRACK hematopoietic differentiation dataset from Singh et al. (2025). **e**, LOTUS-derived lineage-tree representation of STRACK hematopoietic differentiation. The inferred lineage tree integrates velocity-guided edges, growth-weighted branch structure and stage-informed layout, revealing branching trajectories from HSC and MPP states toward erythroid, megakaryocytic, basophil, monocyte and neutrophil fates. **f,** UMAP visualization of the STRACK scRNA-seq hematopoietic differentiation dataset, colored by cell type annotations. **g**, Temporal dynamics of transcription factors along GMP- and MEP-associated differentiation programs in the STRACK dataset. Heatmap values indicate expression z-scores over inferred developmental time. **h**, Volcano plots of GMP- and MEP-associated lineage perturbation screens. Significant genes with positive and negative perturbation effects are highlighted in orange and purple, respectively, and representative regulators are annotated. **i**, Quantification of expected marker and transcription factors recovery in GMP- and MEP-side among top 200 ranked perturbation screens. Bars show the fraction of top-ranked genes corresponding to lineage-associated markers recovered by LOTUS, DeepRUOT or both methods.

To systematically identify genes associated with lineage bias, we performed an *in silico* perturbation screen over the top 2,000 highly variable genes, perturbing each candidate gene in early progenitors and quantifying its effect on predicted neutrophil and monocyte fate probabilities. Predicted perturbation effects showed clear directionality and lineage specificity across the two myeloid fate screens **(Fig. 4b).** In the neutrophil fate screen, perturbations of genes associated with neutrophil differentiation and granule formation increased the predicted Neu fate probability, including *Cebpe*, a regulator of terminal granulocyte maturation, and the canonical neutrophil granule-associated effector genes *Mpo, Elane* and *Ctsg*^40,41^. By contrast, monocyte-associated genes, including *Ccr2, Csf1r, Vcan* and *Klf4*, were located in the negative-effect region, suggesting that enhancement of monocyte programs reduces the predicted probability of differentiation toward the neutrophil fate^42^. Consistent with this pattern, the direction of effects was largely reversed in the monocyte fate screen: *Ccr2, Csf1r, Vcan* and *Klf4* showed positive effects, whereas *Cebpe* and genes associated with other hematopoietic branches, including *Gata2, Cpa3* and *Itga2b*, showed negative effects^11^. Together, these results show that LOTUS-based in silico perturbation screening prioritizes candidate genes consistent with known lineage-associated transcriptional programs and recovers opposing predicted effects of Neu and Mo programs on downstream fate allocation.

To assess whether perturbation rankings recovered expected lineage programs, we quantified the fraction of predefined lineage markers and transcription factors recovered across increasing ranking depths **(Fig. 4c).** LOTUS showed higher recovery of expected genes among the top-ranked candidates in both monocyte and neutrophil screens, with a clearer advantage in the neutrophil screen. These results indicate that lineage constraints improve the biological specificity of perturbation-gene prioritization.

Beyond terminal fate-level perturbation effects, we examined whether the continuous trajectories inferred by LOTUS preserved the temporal dynamics of lineage-associated transcription factors. Along the LARRY differentiation trajectories, distinct hematopoietic branches displayed fate-specific transcription factor programs: neutrophil-associated factors such as *Cebpe* and *Gfi1* increased along the neutrophil branch, monocyte-associated factors such as *Irf8* and *Klf4* increased along the monocyte branch, and megakaryocyte, basophil and mast cell branches showed corresponding lineage-associated transcription factor signatures **(Fig. 4d).** These results indicate that LOTUS-inferred trajectories can organize fate-specific transcription factor programs along branch-resolved differentiation paths.

### LOTUS extends in silico perturbation analysis to an independent hematopoietic dataset

To test whether LOTUS-based perturbation analysis extends across datasets, we applied it to the STRACK hematopoietic lineage-tracing dataset from Singh et al.^43^, which spans HSC/MPP states and erythroid, megakaryocyte, basophil, monocyte and neutrophil fates **(Fig. 4f).** For the present analysis, we restricted the STRACK dataset to wild-type clones and excluded clones carrying induced leukemic driver mutations, thereby focusing on the unperturbed hematopoietic differentiation landscape. The LOTUS-reconstructed lineage tree originated from HSC-enriched regions and connected to multiple mature hematopoietic fates, recapitulating the hematopoietic hierarchy in this independent dataset **(Fig. 4e).** Along LOTUS-inferred trajectories, GMP-associated branches showed elevated expression of myeloid regulators such as Spi1 and Irf8, whereas MEP-associated branches showed increased expression of erythroid and megakaryocytic regulators such as Gata1, Fli1, and Mef2c, indicating recovery of branch-specific regulatory programs **(Fig. 4g).**

In a STRACK in silico perturbation screen over the top 2,972 highly variable genes, myeloid and granulocytic genes such as *Csf3r* and *Ctsg* increased predicted GMP-side fate probability, whereas MEP-associated genes such as *Itga2b, Gata1, Gp5* and *Gp9* showed negative effects. Conversely, *Itga2b* and *Gata1* increased predicted MEP-side fate probability, whereas myeloid genes such as *Spi1, Ctsg* and *Csf3r* reduced it^44,45^ **(Fig. 4h).** These results indicate that LOTUS perturbation predictions capture opposing lineage-regulatory directions between the GMP and MEP branches. Among the top 200 ranked perturbation genes, LOTUS recovered expected lineage markers, including basophil markers in the GMP-side screen and megakaryocyte markers in the MEP-side screen, whereas DeepRUOT introduced non-MEP lineage signals in the MEP-side screen **(Fig. 4i).** Together, these results show that clonal lineage constraints improve the biological specificity of candidate perturbation-gene prioritization.

### LOTUS resolves multi-fate clone potential and dual-lineage regulatory states

We next evaluated whether LOTUS could resolve clone-level multi-fate potential. For representative clones with dual-fate or multi-fate outputs, LOTUS reconstructed continuous differentiation trajectories from day 2 initial cells toward day 6 terminal states, and the generated paths overlapped the experimentally observed day 4 and day 6 clone descendants in state space, indicating that LOTUS can recover differentiation paths through which a single clone contributes to multiple terminal fates **(Fig. 5a).** At the single-cell level, repeated LOTUS simulations from each day 2 initial cell estimated its allocation across hematopoietic terminal fates. The predicted fate composition was concordant with the observed clonal output, with a mean total variation (TV) distance of 0.402 ± 0.014 across 846 evaluable clones (mean ± s.e.m. across clones; Fig. 5a).

**Fig. 5.**
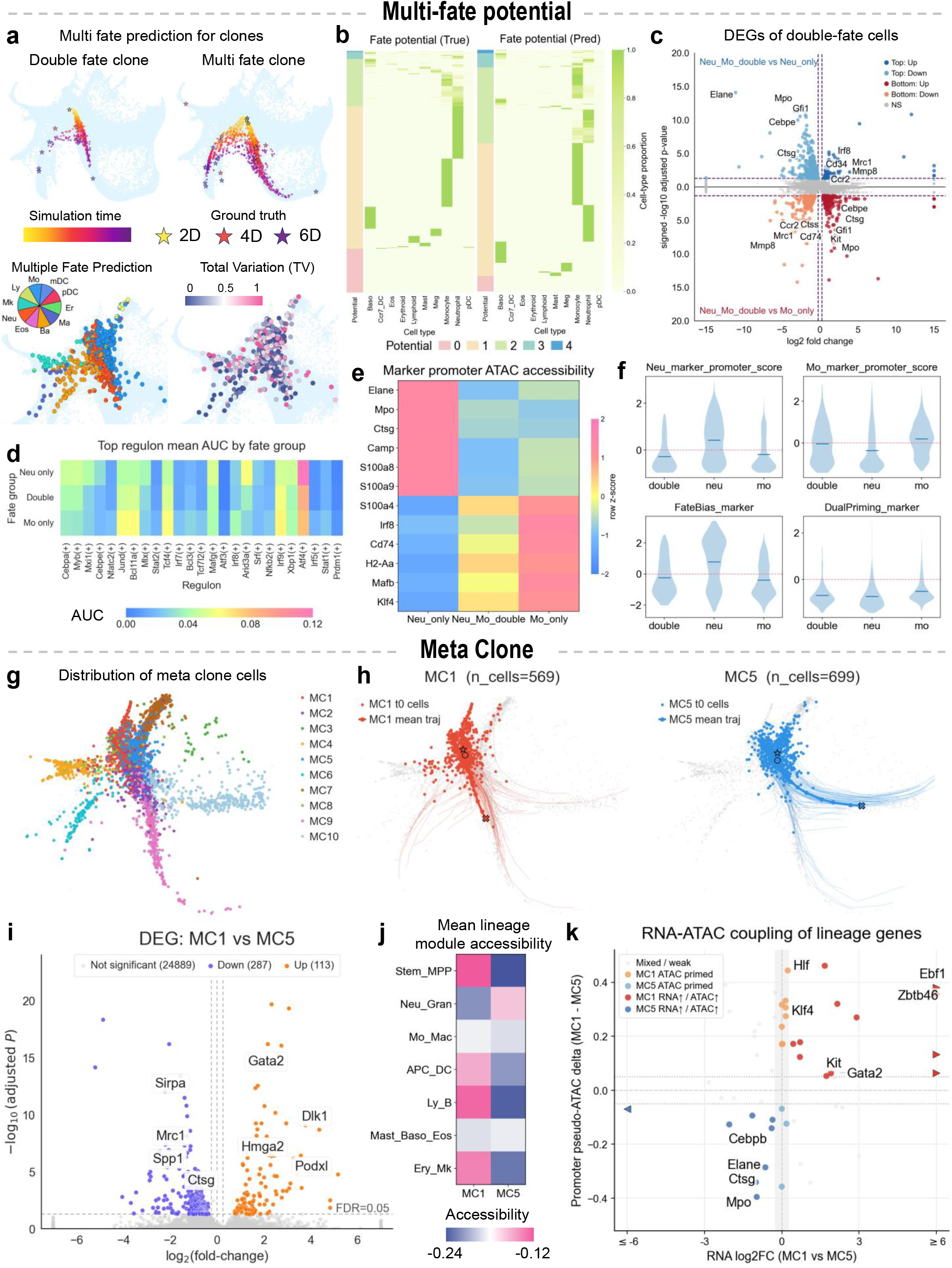
LOTUS resolves multi-fate clone potential, meta-clone identity and clone-associated regulatory programs. **a,** LOTUS-estimated cell differentiation potential. Top, representative double-fate and multi-fate clones are shown across simulated time, with predicted states colored by simulation time and experimentally observed clone states marked at days 2, 4 and 6. Bottom left, predicted fate allocation of a multi-fate clone across terminal hematopoietic states, with the inset pie chart showing the corresponding fate composition. Bottom right, total variation distance between predicted and observed clone fate distributions, highlighting regions where clone-level fate prediction is more or less concordant with the measured lineage output. **b,** Comparison between observed and LOTUS-predicted fate-potential matrices across cell types. Rows represent individual clones or clone groups, and columns denote potential downstream fate states. The leftmost column represents the number of possible differentiation directions for the clone. **c,** Differential expression analysis of Neu-Mo-double-fate cells compared with Neu-only and Mo-only cells. Cells were defined as Neu–Mo double-fate cells if the predicted probabilities for both neutrophil and monocyte fates exceeded 0.05 at any simulated time point. The upper part compares double-fate cells with Neu-only cells, while the lower part compares double fate cells with Mo only cells. Volcano plots highlight genes enriched or depleted in double-fate states relative to single-fate populations. **d,** Mean AUCell activity of differentially active regulons across Neu-only, double-fate and Mo-only cells. The top 25 regulons with differences across the three groups are shown; colors indicate the mean regulon AUC in each group. **e,** Marker promoter accessibility profiles across fate groups, showing differential chromatin accessibility of lineage-associated genes in Neu-only, Neu–Mo double-fate and Mo-only populations. **f,** Distribution of promoter or marker scores for Neu-associated, Mo-associated, fate-bias and dual-priming gene sets across Neu–Mo double-fate, Neu-only and Mo-only groups. **g,** Distribution of meta-clones constructed from predicted clone trajectories in the initial cell embedding space. **h,** Representative trajectories of two meta-clones, MC1 and MC5, indicating that clone-level fate differences can emerge even among cells with highly similar transcriptomic profiles. **i,** Differential expression analysis between MC1 and MC5 cells, identifying clone-associated transcriptional programs. **j,** Mean lineage module accessibility of MC1 and MC5 across major hematopoietic lineage modules. **k,** Coupling between RNA expression changes and promoter inferred-ATAC accessibility for lineage-associated genes. The x axis shows RNA log2 fold change between MC1 and MC5, and the y axis shows promoter inferred-ATAC accessibility difference, highlighting concordant and discordant regulatory programs associated with clone-specific fate bias.

At the global clone level, the LOTUS-predicted fate-potential matrix showed a distribution similar to the experimentally observed matrix, distinguishing single-fate, dual-fate and multi-fate clones **(Fig. 5b).** LOTUS showed better agreement with the observed clone fate distributions than DeepRUOT, with higher TV-derived similarity (0.755 versus 0.664) and normalized Jensen–Shannon divergence (JSD)-derived similarity (0.920 versus 0.892) **(Fig. 5b).** These results support the ability of LOTUS to recover clone-level multi-fate potential across individual clones and the global clone population.

We then focused on Neu–Mo double-fate cells, which were predicted to retain both neutrophil and monocyte potential. Although these cells partially overlapped with single-fate cells in transcriptional space **(Supplementary Fig. 6a),** differential expression analysis showed that Neu–Mo double-fate cells were transcriptionally distinct from both Neu-only and Mo-only cells. Compared with Neu-only cells, double-fate cells showed lower expression of terminal neutrophil-associated genes, including *Elane, Mpo, Cebpe* and *Ctsg*, and relatively higher expression of monocyte-or progenitor-associated genes, including *Irf8, Mrc1* and *Ccr2*. Compared with Mo-only cells, however, double-fate cells retained expression of neutrophil-lineage genes such as *Cebpe, Ctsg, Gfi1* and *Mpo* **(Fig. 5c).** Consistently, gene module scoring showed that Neu– Mo double-fate cells had an intermediate neutrophil score between Neu-only and Mo-only cells while retaining monocyte-associated signal, supporting their interpretation as a Neu/Mo dual-primed state **(Supplementary Fig. 6b,c).**

To assess whether Neu–Mo double-fate cells showed distinct regulatory features, we used SCENIC to compare regulon activity across Neu-only, Neu–Mo double-fate and Mo-only cells^46^. The top 25 differentially active regulons included myeloid-associated regulators such as Cebpa, Cebpb, Irf7, and Irf8 **(Fig. 5d).** Neu– Mo double-fate cells did not simply recapitulate either the Neu-only or Mo-only regulatory profile; instead, they showed intermediate or mixed area under the curve (AUC) activity across multiple Neu-and Mo-associated regulons, consistent with a primed regulatory state that retains features of both lineages.

We next examined putative chromatin-accessibility correlates of these lineage differences using MultiVI-inferred ATAC signals^47^. Paired RNA–ATAC multiome data from ex vivo-expanded murine hematopoietic stem cells and their progeny reported by Zhang et al. were used as the reference dataset for MultiVI inference, such that the inferred accessibility profiles may reflect cell states represented in this reference^48^. Marker promoter accessibility analysis showed that neutrophil-associated genes, including *Elane, Mpo, Ctsg, S100a8* and *S100a9*, had higher inferred accessibility in Neu-only cells, whereas monocyte-associated genes, including *Irf8, Cd74, H2-Aa, Mafb* and *Klf4*, had higher inferred accessibility in Mo-only cells **(Fig. 5e).** By contrast, Neu–Mo double-fate cells retained inferred promoter accessibility at both Neu-and Mo-associated marker loci, displaying accessibility patterns intermediate between the two single-fate populations. Gene-set score analysis further showed that double-fate cells retained both Neu-and Mo-associated regulatory features and were distinguishable from single-fate cells by fate-bias scores **(Fig. 5f).** Together, these results indicate that LOTUS-identified double-fate cells show interpretable molecular and regulatory features, making them unlikely to reflect prediction noise alone.

### LOTUS resolves hidden meta-clone fate biases among transcriptionally similar progenitors

Meta-clones aggregate individual clones with similar fate trajectories or dynamical behaviors into higher-order clonal units, and clones within the same meta-clone are expected to share related differentiation trajectories and fate biases, potentially reflecting common early lineage priming^30^. Meta-clone analysis mitigates clone-level sparsity caused by small clone sizes and provides a stable unit for identifying shared fate biases and regulatory programs among transcriptionally similar early cells. Using LOTUS-predicted clone trajectories, we partitioned early cells into ten meta-clones **(Fig. 5g and Supplementary Fig. 7a),** among which we identified two similarly sized meta-clones, MC1 (n = 569 cells) and MC5 (n = 699 cells). Although MC1 and MC5 partially overlapped in the initial transcriptional embedding, their LOTUS-inferred trajectories showed distinct fate biases: MC1 showed broader downstream fate potential, whereas MC5 was primarily biased toward neutrophil and monocyte lineages. These results suggest that early cells with similar transcriptional states can nevertheless carry distinct state–fate landscapes and clone-associated fate biases **(Fig. 5h).**

We next compared the transcriptional profiles of MC1 and MC5 to identify molecular programs associated with meta-clone-specific fate bias. Differential expression analysis showed that MC1 cells upregulated genes associated with early progenitor or stem-like states and broader non-granulocytic fate potential, including Gata2 and Hmga2. By contrast, MC5 was enriched for myeloid differentiation-associated genes, including *Mrc1, Spp1* and *Ctsg*, consistent with the distinct differentiation trajectories of these two meta-clones **(Fig. 5i).**

To investigate putative chromatin-accessibility correlates of meta-clone-specific fate bias, we compared lineage module accessibility between MC1 and MC5 using MultiVI-inferred ATAC signals. The two meta-clones showed systematic differences in inferred lineage-associated chromatin accessibility **(Supplementary Fig. 7d).** Consistent with its broader inferred trajectories (Fig. 5h), MC1 showed higher inferred accessibility in the Stem/MPP module and several alternative lineage modules, including APC/DC, Ly/B and Ery/Mk, suggesting that MC1 retains a broader, putatively multi-lineage-primed accessibility state. By contrast, MC5, which was primarily biased toward neutrophil and monocyte lineages, showed higher inferred accessibility in Neu/Gran-related modules and lower inferred accessibility in alternative lineage modules **(Fig. 5j).** Consistently, the commitment score, reflecting the degree of bias toward a single lineage program, was significantly lower in MC1 than in MC5. **(Supplementary Fig. 7c).** These results suggest that the divergent trajectories of MC1 and MC5 are accompanied by distinct inferred chromatin-accessibility states.

RNA–ATAC coupling analysis further linked meta-clone fate bias to coordinated, and partially uncoupled, RNA–accessibility patterns **(Fig. 5k).** For a subset of genes, RNA expression changes were concordant with inferred promoter ATAC differences. On the MC1 side, concordant increases in *Kit, Gata2, Ebf1* and *Zbtb46* were consistent with progenitor, lymphoid/B cell and dendritic-cell-related priming^49^. On the MC5 side, concordant increases in *Elane, Ctsg* and *Mpo* reflected granulocytic differentiation programs, whereas *Cebpb* was consistent with monocyte/myeloid inflammatory programs. These results indicate that some of the lineage-associated chromatin-accessibility differences between MC1 and MC5 were accompanied by detectable transcriptional differences. At the same time, genes such as *Hlf* and *Klf4* showed promoter accessibility bias but weaker RNA expression differences, suggesting putative chromatin-primed regulatory features not yet accompanied by strong transcriptional activation. Thus, meta-clone-specific fate bias is reflected in both active transcriptional programs and putative chromatin-primed features.

### LOTUS reconstructs lineage-resolved reprogramming trajectories and fate-associated regulatory programs

To test whether LOTUS extends beyond hematopoietic differentiation, we applied it to the CellTagging reprogramming scRNA-seq dataset from Biddy et al.^22^, which captures multi-time-point state transitions during fibroblast reprogramming. In the UMAP embedding, reprogrammed and failed cells occupied largely distinct terminal regions **(Fig. 6b).** The LOTUS-inferred lineage tree originated from early cell states and branched along velocity-supported directions toward reprogrammed and failed terminal outcomes, recapitulating the major fate structure of the reprogramming process **(Fig. 6a).**

**Fig. 6.**
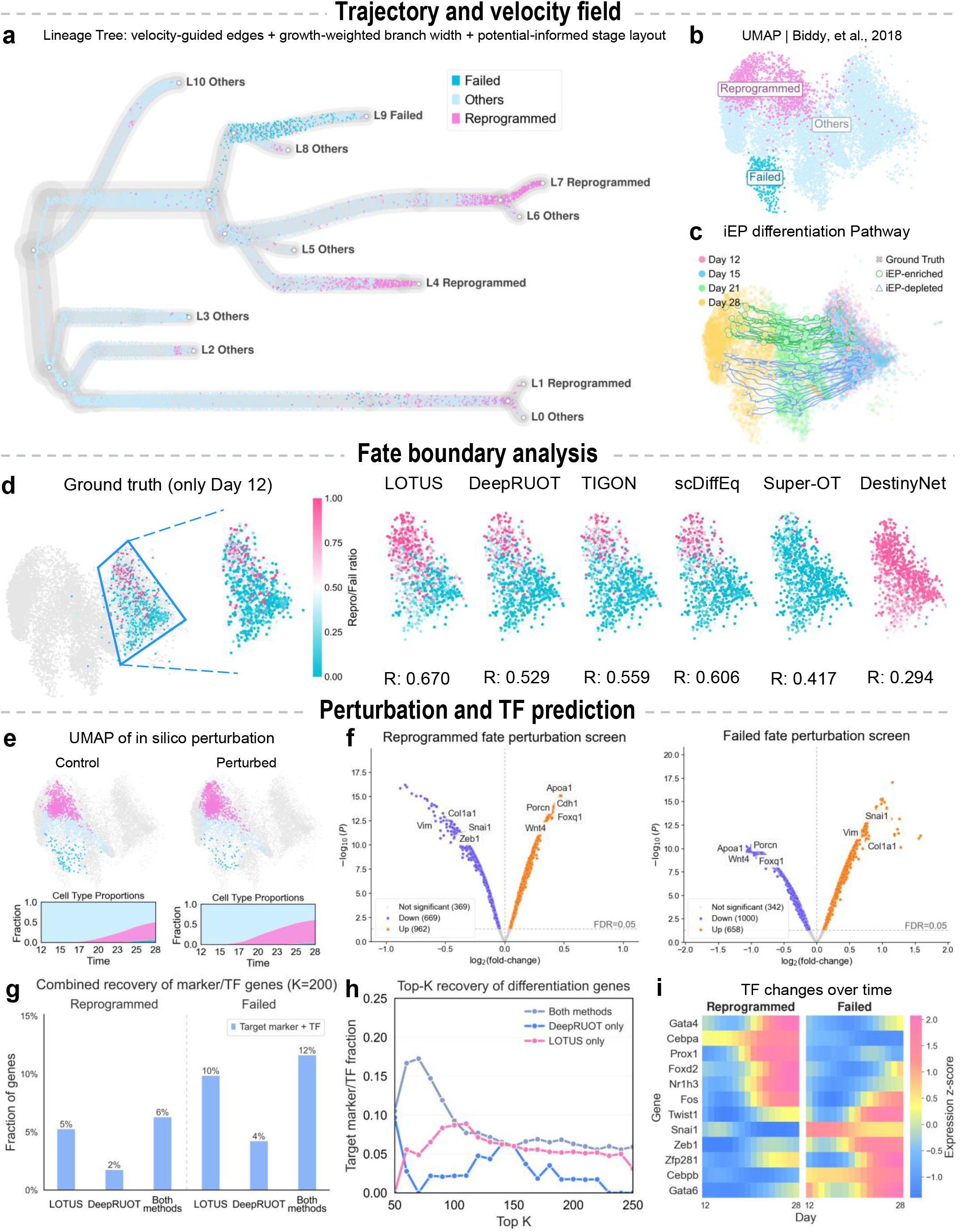
LOTUS reconstructs lineage-dependent reprogramming dynamics. **a**, Lineage tree inferred for the Biddy et al. reprogramming dataset. Edges are guided by local velocity, branch width reflects inferred growth, and the layout is arranged according to developmental potential. Cells are colored by experimentally annotated outcomes. **b**, UMAP visualization of the reprogramming dataset, showing the global organization of reprogrammed, failed and other cell states. **c**, Simulated differentiation pathways for iEP-enriched and iEP-depleted initial populations. Ground-truth cells are shown together with inferred trajectories across days 12, 15, 21 and 28, illustrating divergence between reprogramming-prone and reprogramming-resistant routes. **d,** Fate-boundary prediction for reprogrammed and failed fates. Ground-truth reprogrammed-to-failed fate ratios for Day 12 cells and predictions by LOTUS, DeepRUOT, TIGON, scDiffEq, Super-OT and DestinyNet are shown for the highlighted region. Colors indicate the reprogrammed-to-failed fate ratio, and Pearson correlation *R* quantify agreement with the ground truth. **e,** *In silico* perturbation of the reprogramming landscape after Foxd2 perturbation. Control and Foxd2-perturbed UMAPs show changes in final cell-state distributions, with stacked area plots summarizing the corresponding temporal shifts in cell-type proportions. **f**, Volcano plots showing genes whose perturbation is predicted to promote reprogrammed or failed fates. Genes with significant positive and negative effects are highlighted, and representative regulators are annotated. Dashed lines indicate the FDR threshold. **g**, Recovery of known marker and transcription-factor genes among top-ranked perturbation candidates for reprogrammed and failed fate screens. Bars compare genes recovered by LOTUS, DeepRUOT and genes shared by both methods. **h,** Top K recovery of differentiation-associated genes in the reprogramming screen. The y axis shows the fraction of target markers or transcription factors recovered among the top-ranked candidate genes. Genes are grouped according to whether they were identified by both methods, DeepRUOT only or LOTUS only. **i,** Temporal dynamics of transcription factors associated with reprogrammed and failed outcomes. Heatmap values represent gene-wise expression z-scores over inferred reprogramming time, highlighting regulators with fate-specific temporal activation patterns.

We next asked whether LOTUS could resolve differences in early reprogramming potential. Starting from experimentally defined induced endoderm progenitor (iEP)-enriched and iEP-depleted initial populations, LOTUS generated divergent trajectories: iEP-enriched cells were preferentially directed toward the reprogrammed fate, whereas iEP-depleted cells were biased toward failed or other states **(Fig. 6c).** Consistently, the LOTUS-inferred fate boundary recapitulated the reference reprogrammed-to-failed fate ratio assigned to early day 12 cells from their later clonal outputs, achieving the highest correlation with the reference (*R* = 0.670), outperforming scDiffEq (*R* = 0.606), TIGON (*R* = 0.559), DeepRUOT (*R* = 0.529), Super-OT (*R* = 0.417) and DestinyNet (*R* = 0.294; **Fig. 6d**).

We then examined whether LOTUS could support in silico perturbation analysis during reprogramming. As a proof of concept, Foxd2-associated perturbation increased the predicted bias toward the reprogrammed region **(Fig. 6e).** In highly variable gene-level perturbation screens, LOTUS prioritized directionally coherent candidate genes. In the reprogrammed-fate screen, epithelial or iEP-associated genes, including *Apoa1, Cdh1, Foxq1, Porcn* and *Wnt4,* showed positive predicted effects^50,51^, whereas mesenchymal, EMT-like or extracellular-matrix-associated genes, including *Vim, Col1a1, Snai1* and *Zeb1*, decreased the predicted probability of reprogramming^52,53^. Conversely, these effect directions were largely reversed in the failed-fate screen **(Fig. 6f),** consistent with an antagonistic axis between epithelial-like reprogramming and fibroblast-like failed states.

Top-ranked perturbation genes further recovered known fate-associated markers and transcription factors. Among the top 200 ranked genes and across top-K cutoffs, LOTUS recovered reprogramming-associated marker and transcription factor programs more consistently than DeepRUOT in both reprogrammed-and failed-fate screens **(Fig. 6g,h).** Along the LOTUS-inferred reprogramming axis, *Gata4, Cebpa* and *Foxd2* increased along the reprogrammed route, whereas Snai1 and Zeb1 were enriched along the failed route, consistent with branch-specific epithelial/iEP-like and mesenchymal/EMT-like programs **(Fig. 6i).** Together, these results show that LOTUS reconstructs lineage-resolved reprogramming trajectories and prioritizes fate-associated perturbation genes with coherent branch-specific regulatory dynamics.

### LOTUS reconstructs clonotype-resolved TR1 dynamics during malaria infection

To test whether LOTUS can use endogenous TCR-derived clonal identities for lineage-aware dynamical modeling, we applied it to a longitudinal scRNA-seq and TCR-seq dataset of 504,041 human memory (CD45RA-) CD4+ T cells profiled across malaria infection and stimulation conditions^54^. Expanded resting clonotypes were grouped into seven clone families according to subset preference, and shared TCR sequences linked resting clonotypes to their activated counterparts, providing a reference for tracking clone-family identities across activation states.

Using this TCR-based clone-tracking reference, LOTUS reconstructed activated-state trajectories that maintained the resting-defined clone-family structure **(Fig. 7a and Supplementary Fig. 10a).** Reconstructed cells showed broadly concordant clone-family expression patterns with observed cells, both for family 7 and across global memory T cell subset markers after TCR activation **(Fig. 7b,c).** Concordant MYC-associated regulon activity was also observed in reconstructed activated states **(Supplementary Fig. 10b).**

**Fig. 7.**
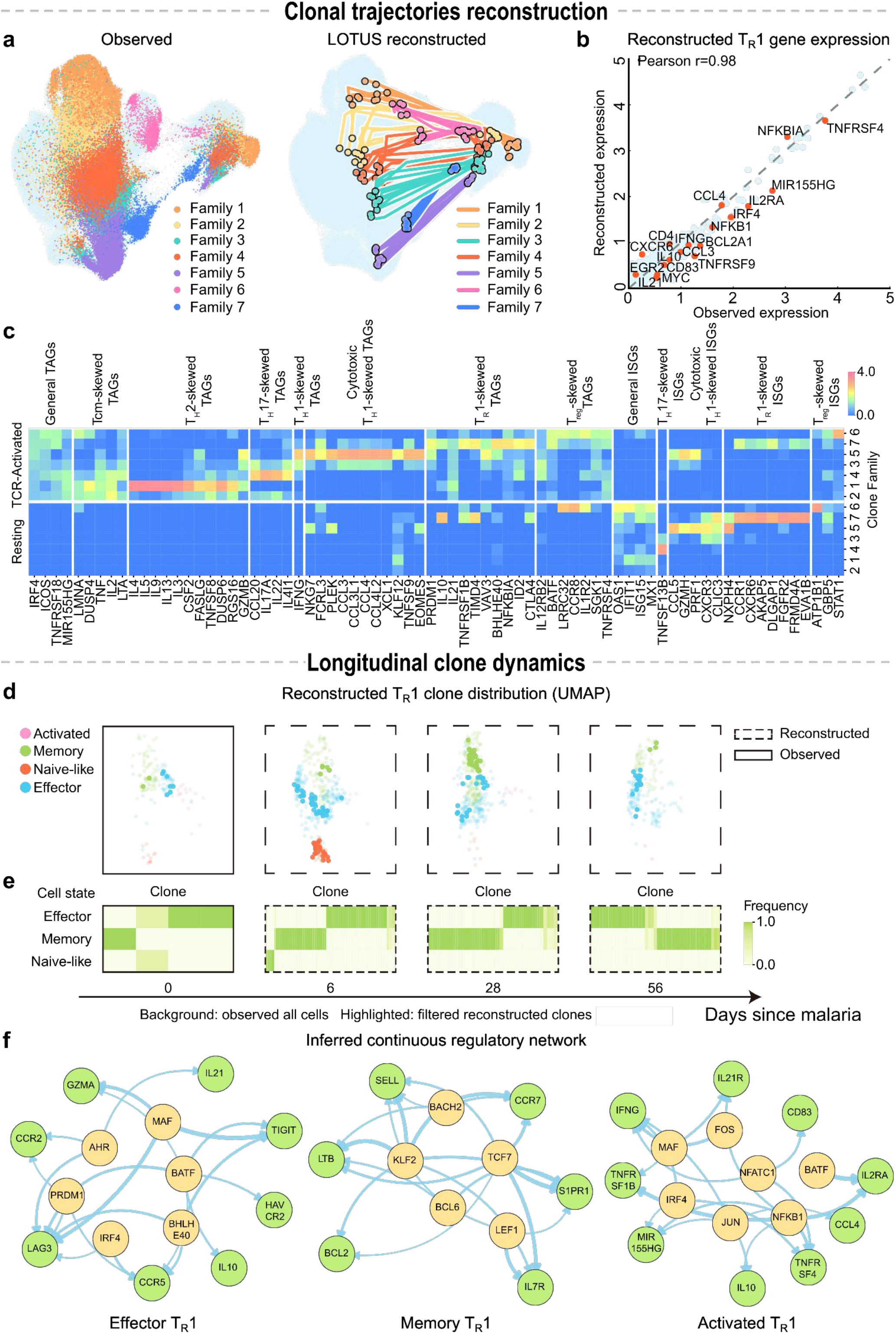
LOTUS reconstructs clonal identity and T_R_1 dynamics in malaria-responsive CD4+ T cells. **a,** Observed clone-family distribution and LOTUS-inferred trajectories. Left, observed CD4+ T cells projected in UMAP space, with colors indicating clone families 1–7. Right, LOTUS-inferred clonal trajectories originating from 10 sampled cells per clone family, projected onto the same embedding; colors denote clone-family identity. **b,** Observed and LOTUS-reconstructed gene expression in family 7 cells. Mean log-normalized expression was compared between observed and LOTUS-reconstructed cells at the matched time point. Highlighted genes denote representative T_R_1-associated differentially expressed genes. Pearson’s r = 0.98 reflects in-sample agreement at an observed training time point, not independent validation. **c,** LOTUS-reconstructed gene expression across clone families. Heatmap showing genes broadly expressed across CD4+ T cells or selectively enriched in specific clone families following TCR activation. Expression patterns were reconstructed by LOTUS. **d,** Reconstruction of longitudinal TR1 clonal dynamics in participant 3481 during malaria infection. Background points indicate the observed distribution of all TR1 cells at each time point, whereas highlight endpoints represent LOTUS-reconstructed descendant cells. **e,** Predicted clonal state composition of TR1 clonotypes from clonotypes containing at least two cells. Each heatmap is vertically aligned with the corresponding UMAP panel above. Columns represent individual clonotypes, rows represent TR1 cell states, and color intensity indicates the proportion of cells in each clonotype occupying the corresponding state. **f,** Scribe-inferred TF-to-gene temporal regulatory networks derived from LOTUS-reconstructed effector T_R_1 (left), memory T_R_1 (middle), and activated T_R_1 (right) dynamics. Yellow nodes represent transcription factors, green nodes represent genes, and edge width is proportional to the inferred temporal dependency strength (cRDI score).

We next applied LOTUS to malaria-responsive TR1 cells from participant 3481, who was extensively sampled during a single symptomatic malaria episode. To focus on the acute post-infection phase, we restricted the trajectory analysis to samples collected on days 0, 6, 28, and 56. LOTUS reconstructed continuous trajectories spanning activated, naïve-like, effector and memory T_R_1 states, consistent with the reported early enrichment of effector T_R_1 cells followed by later contraction **(Fig. 7d and Supplementary Fig. 10c,d).** At clonotype resolution, predicted T_R_1 state compositions linked expanded effector and memory TR1 states through shared clonotype structure **(Fig. 7e).** These results extend discrete longitudinal observations into a continuous view of malaria-responsive TR1 dynamics.

Scribe^55^ analysis of LOTUS-derived trajectories identified state-associated regulatory networks **(Fig. 7f):** effector-associated networks were enriched for T_R_1 regulatory factors such as *BATF, MAF* and *PRDM1*, memory-associated networks highlighted *TCF7, LEF1* and *BACH2*, and activated T_R_1 networks involved TCR-responsive NF-κB, NFAT and AP-1 programs^56,57^.

## Discussion

Time-resolved lineage-tracing single-cell sequencing links transcriptional state, sampling time and clonal identity, providing a basis for reconstructing how cell-state transitions are coupled to clonal fate inheritance. To address the missing ancestor–descendant links inherent to destructive single-cell sampling, we developed LOTUS, a lineage-informed neural optimal transport framework that reconstructs continuous cell-state dynamics while modeling stochasticity and non-conserved changes in cellular mass. LOTUS incorporates lineage barcodes as clone-level constraints within a neural unbalanced optimal transport objective, requiring predicted descendants to match both the global transcriptomic distribution and the observed clone-specific distributions at later time points. The learned dynamics support fate-boundary inference, clone-level fate-potential analysis and in silico perturbation-based gene prioritization.

LOTUS bridges two previously separate modeling strategies for time-resolved single-cell data. Lineage-aware methods such as CoSpar^24^, moslin^28^, LineageOT^27^ and CLADES^30^ use clonal information to infer fate bias, cross-time couplings or clone-level population dynamics, but they generally represent fate dynamics as transition maps, couplings or population-level summaries. Conversely, continuous generative models such as TIGON^32^, DeepRUOT^33,34^ and scDiffEq^35^ learn trajectories from time-series single-cell data but do not directly constrain those trajectories by observed clonal propagation. By combining within-clone optimal transport with neural unbalanced transport dynamics, LOTUS integrates these two principles into a single framework for modeling continuous transcriptomic change, non-conserved cellular mass and lineage-consistent clonal propagation.

Across simulated benchmarks and biological lineage-resolved datasets, LOTUS showed that clonal information can improve the lineage consistency of continuous fate reconstruction. In bifurcating simulations, clone-level supervision reduced off-target routing under increasing clonal complexity, indicating that lineage constraints can resolve transport choices that are ambiguous on the basis of transcriptomic geometry alone. In the LARRY hematopoietic dataset^11^, LOTUS reconstructed continuous myeloid differentiation trajectories, improved neutrophil–monocyte fate-composition prediction against clone-derived reference outcomes and prioritized perturbation candidates consistent with known lineage programs. Its application to STRACK hematopoiesis^43^, fibroblast reprogramming^22^ and clonotype-resolved CD4+ T cell responses^54^ further supported the applicability of LOTUS across distinct lineage-resolved systems.

Despite providing a unified lineage-aware dynamical model, LOTUS has several limitations and opportunities for future development. First, LOTUS relies on high-quality lineage barcode information: barcode dropout, barcode homoplasy, small clone sizes or insufficient clone sharing across time points may weaken the constraint imposed by the clone-level loss. In addition, the current clone loss is weighted by the number of cells from each clone represented in the training mini-batch, which may give more highly represented clones greater influence during training while reducing the contribution of rare clones. Future work could incorporate clone-balanced sampling, hierarchical weighting or adaptive loss functions that account for lineage-label confidence. Second, LOTUS relies on numerical integration of neural network-parameterized dynamics, which introduces computational overhead for large-scale datasets; simulation-free or flow-matching approaches may improve scalability^58–60^. Third, reconstruction accuracy depends on temporal sampling design: long intervals between adjacent time points can increase uncertainty in latent paths and expansion histories, whereas overly short intervals may lack identifiable fate-branching signals. Finally, LOTUS currently reconstructs lineage-resolved cell-fate dynamics mainly in transcriptomic state space, and its ability to model epigenomic regulatory states and true perturbation responses remains limited; extending LOTUS to integrate scRNA-seq, scATAC-seq and Perturb-seq data could enable more direct modeling of the multi-layered regulatory mechanisms underlying cell-fate decisions^58,61–63^.

Overall, LOTUS provides a lineage-constrained generative framework for modeling continuous cell-state dynamics from time-resolved lineage-tracing single-cell data. By using observed clonal propagation as a training constraint, LOTUS reconstructs developmental trajectories that are more consistent with lineage history and supports downstream analyses including fate-boundary inference, multi-fate clone analysis and in silico perturbation-based regulator prioritization. More broadly, LOTUS illustrates how clonal ancestry can be converted from an observational label into a dynamical constraint for reconstructing interpretable models of cell-fate decisions.

## Methods

### Data and problem formulation

We considered time-series scRNA-seq data collected at *T*discrete time points *t*_0_ < *t*_1_ < ⋯ < *t_T_*_−1_. At each time point *t_k_*, the data consist of *N_k_*cells represented by expression or low-dimensional state vectors 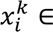 ∈ R*^d^*, giving 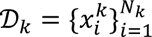. Because scRNA-seq measurements are destructive, cells observed at different time points are unpaired and represent cross-sectional samples from an evolving population.

When lineage tracing information was available, each cell was additionally assigned a clonal barcode 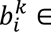, where cells sharing the same barcode were considered descendants of a common labeled progenitor. Cells belonging to uninformative or filtered clones could still contribute to the non-lineage components of the model. Thus, LOTUS does not require every cell to carry a barcode or every observed clone to be used. Although the framework permits unbarcoded cells to contribute to the non-lineage losses, for the analyses reported here we retained only cells with valid lineage barcodes. The resulting data were denoted as D*_k_* = 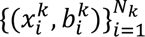, where 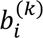 may be missing. These barcode annotations provide lineage constraints that are complementary to transcriptomic similarity.

Given these time-resolved, barcode-labeled snapshots, LOTUS aims to infer continuous cell-state dynamics together with non-conserved changes in cellular mass. We formulated this task as a lineage-aware regularized unbalanced optimal transport problem, in which transport between consecutive time points is constrained by both transcriptomic geometry and observed clonal relationships.

### Data preprocessing

Cells without valid lineage barcodes were removed before analysis. Genes detected in fewer than ten cells were filtered out. Count matrices were normalized to 10,000 counts per cell, log-transformed and restricted to the 2,000 most highly variable genes using Scanpy^64^. The resulting expression matrix was scaled, with values clipped at 10, and reduced to 50 principal components. Time-point annotations were converted to numerical model-time coordinates, with the earliest observation assigned to zero. For the LARRY dataset, days 2, 4 and 6 were encoded as model times 0, 1 and 2, respectively, so that consecutive observation intervals had unit length. The mapping from experimental sampling times to model time was dataset-specific. The PCA coordinates and sample indices were used for model pretraining. Lineage barcodes were introduced during the lineage-aware refinement stage. Precomputed two-dimensional embeddings were used only for visualization. For the STRACK dataset, 2,972 HVGs was used, and batch effects were corrected using Harmony on the 50-dimensional PCA representation with the batch annotation as the integration covariate. The Harmony-corrected PCA coordinates were subsequently used as the cell-state representation for model training and downstream analyses.

### Lineage-aware regularized unbalanced optimal transport

#### Theoretical foundation

We modeled the evolution of the cell population density *p*(**x**, *t*) in gene expression space as a stochastic process governed by the stochastic differential equation (SDE):

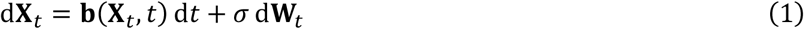

where **b**(**x**, *t*) ∈ R*^d^* is the drift field encoding deterministic cell-state transitions, *σ* > 0 is a constant diffusion coefficient capturing stochastic gene expression fluctuations, and **W***_t_* is a standard *d*-dimensional Brownian motion. The corresponding generally unnormalized cell-population density *p*(**x**, *t*) satisfies the Fokker– Planck equation with a source term accounting for effective changes in cellular abundance.

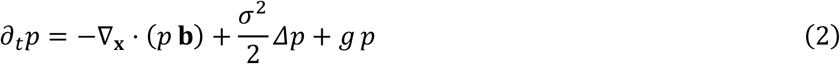

where *g*(**x**, *t*) ∈ R is the local growth and death rate, with positive values indicating net proliferation and negative values indicating net apoptosis.

The regularized unbalanced optimal transport (RUOT) framework seeks to find the velocity field **b**, growth rate *g*, and density *p* that together minimize a total action functional comprising a kinetic energy (quantifying the transport cost) and a growth penalty (regularizing the growth and death process):

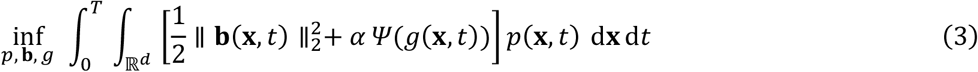

subject to the Fokker–Planck constraint (Eq. 2) and the boundary conditions *p*(⋅, *t*_0_) = *ν*_0_, *p*(⋅, *t_T_*_−1_) = *ν_T_*_−1_, where *ν_k_* denotes the empirical density at time *t_k_*, *α* > 0 controls the strength of the growth penalty, and *Ψ*(*g*) = |*g*|^2^ is the quadratic penalty function. When *g* ≡ 0, the problem reduces to the balanced Schrödinger bridge. When *σ* → 0, it reduces to the unbalanced dynamical optimal transport with the Wasserstein–Fisher–Rao metric.

#### Fisher information regularization and probability flow ODE

Direct computation of the RUOT objective (Eq. 3) is intractable in high dimensions because it requires evaluating the density *p*(**x**, *t*) over the entire state space. Following the Fisher information regularization approach, we reformulated the problem in terms of a probability flow ODE. Specifically, the SDE above can be equivalently described by an ODE that generates the same time-marginal densities:

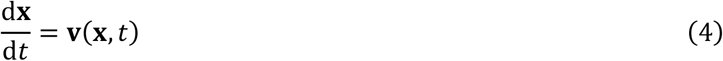

where 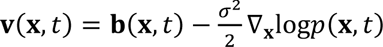 is the probability flow velocity field, which absorbs the score function ∇**_x_**log*p*. Under this transformation, the density evolves according to the continuity equation with source:

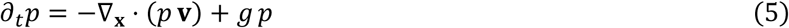

For fixed endpoint densities, the RUOT action can be equivalently reformulated, up to a constant endpoint entropy term, in terms of the probability-flow velocity, growth rate and score function as:

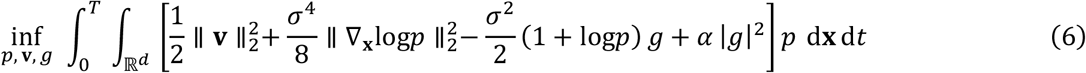

This formulation is computationally advantageous because forward integration of the probability flow ODE (Eq. 4) transforms the high-dimensional spatial integral into a tractable Monte Carlo expectation over initial conditions:

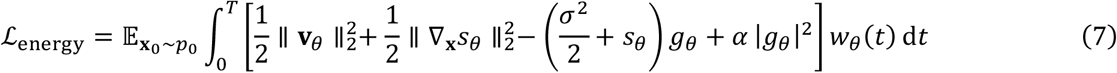

where 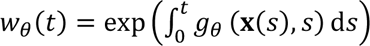 is the cumulative growth factor along each trajectory, and subscripts *θ* denote neural network approximations (see below). In the numerical implementation, we used the Fisher-regularized action together with soft marginal-matching losses, rather than imposing exact endpoint constraints. Consequently, the implemented objective is not strictly equivalent to the original RUOT action under soft endpoint constraints.

### Neural network parameterization

We parameterized three quantities using fully connected neural networks. The velocity network **v***_θ_*(**x**, *t*): R*^d^* × R → R*^d^* represented the probability flow velocity field. The growth network *g_θ_*(**x**, *t*): R*^d^* × R → R represented the local growth and death rate. The scaled log-density network *s_θ_* (**x**, *t*): R*^d^* × R → R approximated 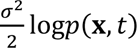, from which the vector-valued score function was obtained through spatial differentiation 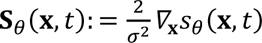.

Given an initial batch of *N* cells 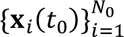 sampled from time *t*_0_, we integrated the coupled ODE system forward in time using a neural ODE solver. The system comprised three components: the trajectory equation d**x***_i_*/d*t* = **v***_θ_*(**x***_i_*, *t*), the weight equation dlog*w_i_*/d*t* = *g_θ_*(**x***_i_*, *t*) with initial condition *w_i_*(*t*_0_) = 1/*N*_0_, and an energy accumulator 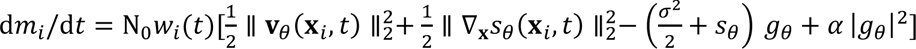 with *m_i_*(t_0_) = 0. The energy accumulator integrates the pointwise energy density weighted by the cumulative growth factor N_0_*w_i_*(*t*), thereby accounting for changes in particle mass along each trajectory. Integration of this system from *t_k_* to *t_k_*_+1_ yielded predicted cell positions **x̂***_i_*(*t_k_*_+1_), updated weights *w_i_*(*t_k_*_+1_), and the accumulated energy cost *m_i_*(*t_k_*_+1_) for each particle.

#### Training objective

During lineage-aware refinement, the training objective comprised four terms that jointly enforced distributional matching, mass consistency, dynamical regularity, and lineage fidelity:

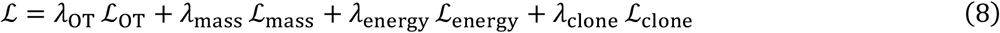

We describe each term below.

#### Optimal transport reconstruction loss

The distributional matching term measured the discrepancy between the predicted and observed cell distributions at each time point. Let 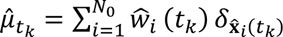 denote the predicted distribution, where 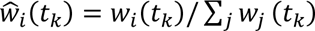 are the normalized predicted weights, and let 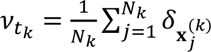 denote the observed distribution with uniform weights. During the initial dynamics-pretraining pass, distributional reconstruction was evaluated using an EMD-based optimal transport loss. During subsequent dynamics refinement and lineage-aware training, we used the differentiable Sinkhorn divergence with a squared Euclidean ground cost:

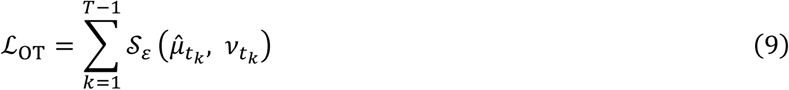

where S_s_ denotes the entropy-regularized Sinkhorn divergence with squared Euclidean ground cost and entropic blur parameter *ε*.

#### Mass matching loss

To encourage the learned growth rates to remain consistent with relative sample-size changes across observation times, we designed a mass-matching strategy that operated at both local and, when enabled, global levels. For each time point *t_k_*, we defined a nearest-neighbor assignment ℎ*_k_* that mapped each observed cell 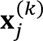 to its closest predicted particle in {**x̂***_i_*(*t_k_*)}:

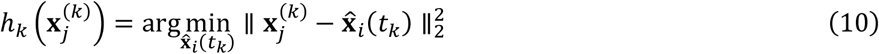

The local mass matching loss penalized discrepancies between the predicted weight of each particle and the fraction of observed cells assigned to it:

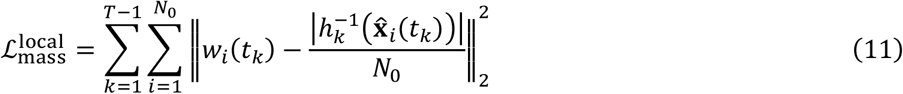

where 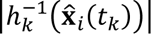 counts the number of observed cells assigned to particle *i*. The local mass-matching term compared the predicted mass associated with each particle with the local abundance of nearby observed cells. When enabled, we additionally imposed an explicit global term that penalized deviations between the total predicted mass and the observed relative sample-size ratio:

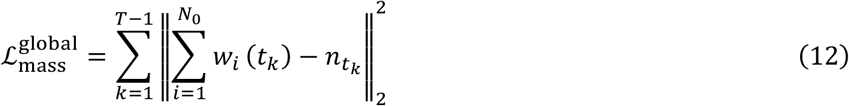

The total mass loss was 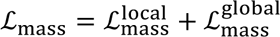. The global term was disabled during the initial pretraining pass and enabled during subsequent dynamics refinement and lineage-aware training.

In mini-batch training, the local target mass was computed by multiplying the fraction of sampled target cells assigned to each predicted particle by the full-data relative cell abundance *N_k_*/*N*_0_, where *N_k_* denotes the number of cells at observation time. Thus, the target masses summed to *N_k_*/*N*_0_ for fixed-size batches, preserving the relative population-mass scale across observation times.

#### Energy regularization

The energy loss implemented the least-action principle of the RUOT functional (Eq. 6), computed as the mean accumulated energy along predicted trajectories:

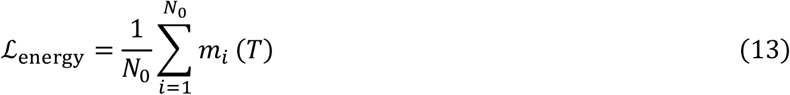

where *m_i_*(*T*) denotes the cumulative energy carried by particle *i*at the end of the current observation interval. The terminal energy value was propagated to the next interval, but detached from the preceding computational graph. This term promoted smooth, low-energy transport paths and penalized excessive velocity and growth contributions according to the action functional.

#### Clonal optimal transport loss

The key methodological advances of our approach was the introduction of a clone-level optimal transport loss that directly exploited lineage barcode information to constrain the inferred dynamics. An initial clone-aware batch was sampled at the first observation time, and each predicted particle retained the barcode inherited from its initial-time ancestor throughout sequential integration. At every subsequent observation time *t_k_*, we identified the barcodes shared between these predicted ancestral lineages and the observed target batch. For each shared clone *c*, we extracted the predicted descendants carrying barcode *c* and the observed target cells bearing the same barcode:

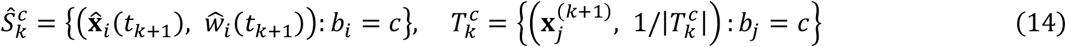

The clone loss was implemented as a mini-batch-count-weighted mean of within-clone debiased Sinkhorn divergences. For each shared clone c, we computed the Sinkhorn divergence between the predicted clone-restricted distribution and the observed clone-restricted target distribution using GeomLoss with a blur parameter of 0.1.

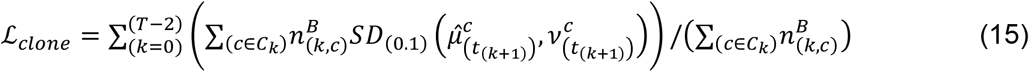

Here, 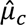 denotes the predicted distribution restricted to particles descended from clone *c*, with growth-derived particle weights normalized within the clone, and *ν_c_*denotes the observed distribution of target cells carrying the same barcode, with uniform target weights. SD_0.1_denotes the debiased Sinkhorn divergence computed using GeomLoss with a blur parameter of 0.1. Per-clone divergences were averaged using the corresponding clone counts in the current mini-batch as weights. Thus, larger clones represented by more cells in the mini-batch contributed proportionally more to the clone loss.

Biologically, this loss term enforced the constraint that cells sharing a common ancestor should be transported to spatial locations consistent with the experimentally observed positions of their clonal relatives at the next time point. This is particularly important when distinct lineages pass through overlapping regions of gene expression space: without lineage information, purely distributional matching cannot distinguish between alternative transport hypotheses, whereas the clonal loss resolves these ambiguities by anchoring the transport map to ground-truth lineage relationships. The clone-level constraint was applied to probability-flow ODE predictions and did not explicitly enforce equivalence between clone-restricted distributions generated by the ODE and the corresponding stochastic SDE.

#### Clone-aware mini-batch sampling

Effective computation of the clonal loss required that cells from the same clone be jointly present in each training mini-batch. Standard uniform random sampling would fragment clonal groups across batches, making the clone loss uninformative. We therefore designed a clone-aware sampling procedure that preserved clonal integrity.

At the initial time point *t*_0_, the sampling procedure first randomly permuted the list of available clones and then sequentially added complete clones until adding another complete clone would exceed the target batch size. If the addition of the final clone exceeded the remaining capacity, a random subset of that clone’s cells was included to exactly fill the batch. The set of clonal barcodes ℬ_0_ present in the resulting batch was recorded for use at subsequent time points.

At each subsequent time point *t_k_*, a priority-based strategy was employed to maximize the overlap of clonal barcodes with the initial batch. All cells belonging to clones in ℬ_0_ were designated as priority cells. When the number of priority cells exceeded *N*, a dispersed sampling scheme was applied: priority clones were randomly ordered, and up to min(nc, m) cells were sampled sequentially from each clone until the batch capacity prevented further allocation, where nc denotes the clone size and m was set to 20 by default. Any remaining capacity was then allocated among priority clones using clone-size-weighted random sampling. When priority cells were insufficient to fill the batch, all priority cells were included and the remaining slots were filled by randomly sampling from non-priority clones. This strategy guaranteed that matching clonal populations were present at both the source and target time points within each mini-batch, enabling meaningful computation of the clone-level transport loss while maintaining batch-size consistency and clonal diversity. The clone-aware source and target mini-batches were also used to evaluate the global transport and mass-matching losses during lineage-aware refinement. Because sampling prioritized shared clones, these losses were evaluated on clone-enriched mini-batch distributions without inverse-probability correction, while the total-mass targets remained calibrated to the full-data cell-count ratios.

#### Score function initialization

The score network *s_θ_* was initialized using conditional flow matching (CFM). Trajectory samples were first generated by integrating the pretrained velocity field forward from the initial time point. Conditional Brownian bridges were then constructed between paired samples at consecutive time points via the optimal transport plan, yielding the conditional density *p*(**x**, *t* ∣ **x**_0_, **x**_1_) = N(**x**; *t* **x**_1_ + (1 − *t*) **x**_0_, *σ*^2^*t*(1 − *t*)). The score network was trained to minimize:

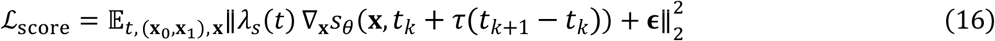

where 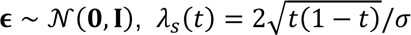 is a time-dependent weighting function, and the expectation was taken over uniformly sampled *t* ∈ [0,1], paired samples from the OT plan, and the conditional Brownian bridge.

The conditional bridges were parameterized using a normalized local interpolation coordinate, while the scaled log-density network received the corresponding global model time. The bridge variance followed a unit-interval convention and was not explicitly rescaled by the duration of each observation interval.

#### Training procedure

We employed a three-stage training strategy consisting of dynamics pretraining, score-network training and lineage-aware refinement. First, the velocity and growth networks were pretrained without lineage information using the optimal transport reconstruction, mass-matching and energy-regularization losses. The pretrained dynamics were subsequently refined for additional epochs with the mass-loss weight unchanged from pretraining (lambda_mass = 0.01) and the energy-loss weight set to zero. Second, trajectories generated by the pretrained dynamics were used to train the score network through conditional flow matching (Eq. 16). Third, lineage information was incorporated through clone-aware mini-batch sampling and the clone-level optimal transport loss. During this lineage-aware refinement stage, the velocity and growth networks were optimized using the full objective in Eq. 8, while the pretrained score network was kept fixed and used to define the probability-flow dynamics. The coupled ODE system was numerically integrated using the Euler method with a fixed step size of 0.1 through the torchdiffeq library. The loss was evaluated and backpropagated separately for each consecutive observation interval. The predicted terminal positions, particle weights and accumulated-energy states were detached from the preceding computational graph and used as the initial conditions for the next interval.

#### Clone-resolved cell-population dynamics simulation

This simulation strategy was used only for the CD4+ T cell dataset. After training, continuous clonal trajectories were generated by forward simulation from the observed initial population using a clone-resolved stochastic cell-population model. Cells from the first observed time point were used as initial particles in the same 50-dimensional PCA state space used for model training. Each particle was initialized with its observed PCA coordinates, assigned an equal initial log-weight, and labeled with its observed lineage barcode. A lineage identifier was also assigned to each initial particle so that descendants generated during simulation could be tracked across time.

For numerical simulation, the learned stochastic cell-population dynamics were integrated on a fixed Euler grid with a step size of 0.1. At each integration step, particle states were first propagated using the learned velocity network *v_θ_* together with the score network *s_θ_*, while particle log-weights *l_i_* were updated using the learned growth network *g_θ_*.

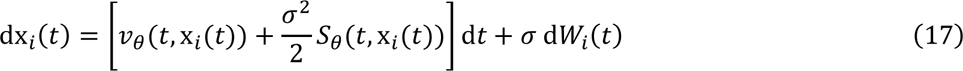

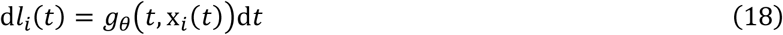

Each particle *i* (*i* = 1, …, *N*) carries a state x*_i_*(*t*) ∈ R*^d^* and a log-weight *l_i_*(*t*) = *logw_i_*(t). In this clone-resolved simulation scheme, cell expansion and loss were evaluated after each Euler update. At time point *t_k_*, the accumulated log-weight change over the previous step was converted into a relative weight ratio:

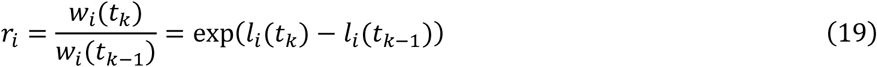

If *r_i_*≥ 1, particle *i* generated a stochastic number of daughter particles:

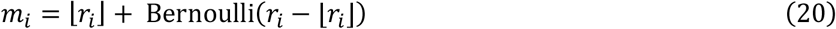

Each daughter particle inherited the updated parent state after the Euler step, with a small additive Gaussian perturbation:

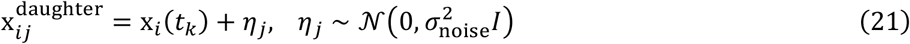

If *r_i_*< 1, particle *i* survived with probability *r_i_* and was removed otherwise. Daughter and surviving particles inherited the parental lineage barcode and lineage identifier, allowing clone-level descendants to be followed across the reconstructed trajectory. After branching or extinction, surviving particles were assigned uniform log-weights relative to the initial particle number:

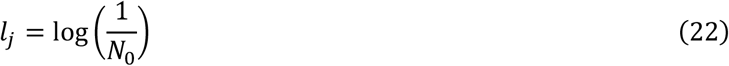

The updated particle population was then used as the input for the next step of the clone-resolved stochastic simulation. In this way, changes in cellular abundance were represented directly by simulated cell counts. The first output time point was fixed to the true observed initial cell population to ensure exact alignment with the empirical initial condition.

### Differentiation potential

To derive a scalar differentiation potential from the trained LOTUS dynamics, we first evaluated the model-implied drift for each observed cell in the 50-dimensional PCA space. The drift was defined by the trained velocity field together with the score-derived correction when diffusion was non-zero, and was subsequently projected onto the two-dimensional UMAP embedding using the velocity-graph framework. We then fitted a neural scalar potential ϕ(u) on the UMAP coordinates u such that its negative gradient approximated the projected drift, V(u) ≈ −∇ϕ(u). The potential network consisted of two hidden layers with 128 units and LeakyReLU activation and was optimized for 2,000 epochs using Adam with a learning rate of 10⁻³ by minimizing the mean squared error ||−∇ϕ(u) − V(u)||².

### Simulated data benchmark

#### Simulated data generation

We generated synthetic time-series datasets with predefined lineage structure to evaluate whether LOTUS could recover lineage-consistent transport when different clonal populations occupied partially overlapping cell-state regions. Each dataset contained two time points and a bifurcating latent manifold with two terminal fate branches.

The ground-truth latent space was defined as a two-dimensional boomerang-shaped bifurcation. Cells were sampled along the two branches using a one-dimensional progression variable, with early cells concentrated near the bifurcation point and later cells distributed farther along the terminal branches. Local cell-state variability was introduced by adding Gaussian noise perpendicular to the branch direction. To create an ambiguous early state, an additional population was sampled near the bifurcation point, where cells from the two lineages partially overlapped in latent space. At the initial time point, 1,500 and 6,000 branch-associated cells were sampled for the two lineages, together with 1,500 cells in a junction-centered mixed population, corresponding to 16.7% of the initial population; at the terminal time point, 6,000 and 12,000 cells were sampled from the two respective branches. The junctional population was sampled from a Gaussian cloud centered at the bifurcation (s.d. = 0.03), whereas progression along each branch was sampled from truncated Gaussian distributions centered at 0.18 (s.d. = 0.18) at the initial time point and 0.72 (s.d. = 0.24) at the terminal time point.

In the two-clone simulation, cells on the upper and lower branches were assigned to two distinct barcodes. The two barcodes differed in both initial abundance and terminal expansion, generating unequal clone sizes and fate-specific growth. This setting was designed to test whether lineage information could disambiguate transport between transcriptionally proximal early cells with different future fates.

We further generated a multi-clone extension using the same latent geometry and sampling procedure. In this setting, each fate branch was divided into multiple clones. Clone identifiers were partitioned into upper-fate (Fate 1) and lower-fate groups (Fate 2), and clone sizes were sampled within each fate group to introduce clonal heterogeneity. Cells in the mixed early population were first assigned to a fate according to their position relative to the bifurcation axis and then assigned to individual clones within that fate in proportion to clone size.

To obtain high-dimensional model input, the two-dimensional latent coordinates were linearly embedded into a ten-dimensional expression-like feature space. The exported synthetic datasets contained the time-point index, ten simulated features and lineage barcode annotations. The original two-dimensional coordinates were retained only for visualization and quantitative evaluation.

#### Benchmark evaluation on simulated data

For the simulated benchmark, LOTUS and all neural-network-based baseline methods were first pretrained on the same randomly generated dataset from the simulation family. The pretrained models were then directly applied to independently generated test datasets without additional training, fine-tuning or parameter updates. Thus, all transferable models were evaluated under the same zero-shot protocol. Because the test datasets shared the same underlying bifurcation geometry, sampling procedure and barcode-to-fate structure, this benchmark assessed whether the dynamics learned from one random realization could generalize to independent realizations of the same data-generating distribution and recover the expected lineage-consistent transport pattern.

For each test dataset, all cells from the initial time point were used as input particles and propagated forward using the pretrained velocity, growth and score networks. The resulting trajectories, terminal predicted states and trajectory weights were recorded. For visualization, principal component analysis was fitted on the observed ten-dimensional simulated features, and the same projection was applied to both predicted states and simulated trajectories. Predicted trajectories were then compared with the observed initial and terminal distributions in the two-dimensional PCA space, with cells colored by time point or lineage barcode.

To quantify reconstruction accuracy, we measured whether each predicted terminal state reached the true terminal neighborhood of its corresponding barcode. For each barcode, observed cells from the final time point were used as terminal reference cells. A local distance score was computed as the mean Euclidean distance from a query point to its 15 nearest terminal reference cells of the same barcode. The true-neighborhood radius for each barcode was estimated from the local distance scores among the observed terminal reference cells of that barcode, excluding self-neighbors. The radius was defined as 1.2 times the maximum reference local score. For each predicted terminal state, the barcode of the corresponding initial cell was used as the expected barcode. A prediction was considered on-target if its local distance score to the terminal reference cells of the expected barcode was smaller than or equal to the barcode-specific true-neighborhood radius. Otherwise, the prediction was considered off-target. The off-target rate was defined as the fraction of predicted terminal states classified as off-target among all propagated initial cells.

#### Baseline comparisons on simulated data. LOTUS was compared with

DeepRUOT, TIGON and scDiffEq using the same ten-dimensional simulated input features and time-point annotations. For all methods, model input consisted of the sample index and the ten simulated expression-like features. Lineage barcode annotations were used only by LOTUS and were not provided to baseline methods unless required for downstream evaluation.

DeepRUOT was run using the DeepRUOTv2 implementation available at https://github.com/zhenyiizhang/DeepRUOTv2. The input dimension was set to 10. The sample size was set to 256, and the score-training batch size was set to 512. All other parameters were kept at the recommended default values provided by the original implementation.

TIGON was run using the implementation available at https://github.com/zhenyiizhang/DeepRUOTv2. The input dimension was set to 10. The sample size was set to 256, and the score-training batch size was set to 512. To obtain deterministic dynamics, sigma was set to 0. All other parameters were kept at the recommended default values provided by the original implementation.

scDiffEq was run using the implementation available at https://github.com/scDiffEq/scDiffEq. Unless otherwise specified, the recommended default parameters were used. The number of training epochs was set to 1,500.

For each method, trajectories were initialized from cells at the first time point and propagated to the terminal time point. Predicted terminal states were projected into the same PCA space as the observed simulated data and evaluated using the terminal true-neighborhood criterion described above. All methods were evaluated on the same independently generated test datasets to ensure a matched comparison of trajectory reconstruction and off-target transport.

### Fate boundary prediction

#### Fate-boundary prediction in the hematopoietic system

To evaluate whether LOTUS could recover early fate bias, we focused on the fate boundary between neutrophil and monocyte outcomes. For each clone, the experimentally observed fate bias at day 2 was inferred using its descendant cells observed at days 4 and 6. Specifically, we calculated the fraction of neutrophil cells among neutrophil and monocyte cells across the clone’s progeny at these later time points to obtain a clone-level Neu/Mo ratio. This ratio was then mapped back to the corresponding undifferentiated cells at day 2 according to their lineage barcode, yielding the reference fate-boundary signal.

For fate-boundary evaluation, the LARRY dataset was divided into two non-overlapping subsets at the clone level, with part 1 used for model training and part 2 used for testing. Each lineage barcode and all cells carrying that barcode were assigned entirely to one subset, ensuring that no clones were shared between the training and test sets. The two subsets were balanced to maintain similar cell numbers, sampling-time distributions and transcriptomic distributions in PCA space.

To obtain the LOTUS-predicted fate boundary, cells collected at day 2, which served as the initial undifferentiated population, were propagated forward to later time points using the trained LOTUS model. Because the generated cell states did not carry cell-type annotations, an auxiliary multilayer perceptron classifier was trained to map continuous cell states to observed cell-type annotations. The classifier contained two hidden layers with 128 units and LeakyReLU activation and was trained for 1,000 epochs using cross-entropy loss and the Adam optimizer. The classifier achieved an overall accuracy of approximately 0.93 and was used solely to assign cell-type labels to model-generated states for downstream fate quantification.

Predicted clone-level fate bias was computed using the same procedure as for the observed data. For each clone, the predicted Neu/Mo ratio was calculated from classified LOTUS-generated cells at later time points and then mapped back to undifferentiated cells at the initial time point by barcode. The agreement between predicted and observed fate-boundary signals was quantified using a clone-size-weighted Pearson correlation across clones. Clones with undefined observed Neu/Mo ratios were excluded, and undefined predicted ratios were assigned a neutral value of 0.5 before correlation calculation. Clone size at the initial time point was used as the weight.

#### Fate boundary prediction in the reprogramming system

For fate-boundary evaluation in the reprogramming dataset, the data were split into training and test subsets using the same general strategy. Owing to the smaller number of cells, clones containing more than 100 cells(n=15) were divided between the two subsets to ensure sufficient sample sizes, whereas all remaining clones were assigned entirely to either the training or test set without overlap. The two subsets were balanced to maintain comparable cell numbers, sampling-time distributions and transcriptomic distributions in PCA space.

Then, the fate-boundary analysis for the reprogramming dataset was performed using the same strategy, except that clone-level fate bias was defined as the fraction of reprogrammed cells among cells assigned to either the reprogrammed or failed state at the final time point and was mapped back to the initial cells by barcode.

#### Fate boundary benchmark

DeepRUOT was run using the DeepRUOTv2 implementation available at https://github.com/zhenyiizhang/DeepRUOTv2. All parameters were kept at the recommended default values provided by the original implementation.

TIGON was run using the implementation available at https://github.com/zhenyiizhang/DeepRUOTv2. To obtain deterministic dynamics, sigma was set to 0, which was the key difference from the DeepRUOT setting. All other parameters were kept at the recommended default values provided by the original implementation.

scDiffEq was run using the implementation available at https://github.com/scDiffEq/scDiffEq. We followed the standard training workflow and fitted a drift–diffusion stochastic differential equation to the precomputed principal-component representation. We set time_key=“samples”, use_key=“X_pca”, kNN_key=“X_pca”, data_dim=50, latent_dim=50, reduce_dimensions=False and num_workers=0, and trained the model for 1,500 epochs using model.fit(train_epochs=1500). All other drift, diffusion and training hyperparameters were retained at their package defaults.

Super-OT was run using the implementation available at https://github.com/uhlerlab/superot. We used the fully supervised Super-OT mode in ‘main.py’, so the model was trained with paired source-target MSE supervision plus identity regularization and no discriminator loss. Super-OT was trained on the clone-holdout CSV splits using the precomputed 50-dimensional PCA features, and the generator input dimension was therefore set to 50 instead of the default nz=100. For multi-time-point datasets, separate generators were trained from the earliest observed time point to each later time point using same-clone source–target pairs. Models were trained for 20,000 iterations. All other supervised training settings, including the generator hidden dimension, batch size, learning rate and identity-regularization weight, were retained at their official default values.

DestinyNet was run using the implementation available at https://github.com/jiangsongtao/DestinyNet. Relative to the official training workflow, we adapted the input format and supervision construction to the clone-holdout benchmark setting. DestinyNet was trained directly from the benchmark CSV splits using the precomputed 50-dimensional PCA features, and len_geneExp was therefore set to 50 instead of the full gene-expression dimension used in the original implementation. For each source cell at the earliest observed time point, one target cell with the same lineage barcode was sampled at each later time point, and a separate mapper was trained for each source-to-target time interval. Pair-classification labels were defined by the target cell types. The number of training epochs was increased from the default 5 to 200. All other settings were retained.

CoSpar was run using the implementation available at https://github.com/AllonKleinLab/cospar. CoSpar was included as a lineage-aware transition-map baseline for fate-boundary prediction, because it infers transition probabilities by integrating transcriptomic similarity with clonal information. CoSpar was evaluated only in the full-dataset setting and was not applied to the train–test split setting, as its transition-map inference requires sufficient shared clonal information across time points.

In the hematopoietic fate-boundary benchmark, cells without valid lineage barcodes were removed before analysis. A transition map was inferred using multi-time-point clonal information, with clonal time points set to days 2, 4 and 6, the later time point set to day 6, smooth_array set to [20, 15, 10], sparsity_threshold set to 0.2 and max_iter_N set to 3. The inferred transition map was restricted to transitions from day 2 cells to day 6 cells. Each row of the restricted transition matrix was normalized to sum to one, and transition probabilities to day 6 neutrophil and monocyte cells were aggregated according to the observed day 6 cell-type labels. The resulting clone-level Neu/Mo ratios were mapped back to day 2 undifferentiated cells by barcode and evaluated using the same clone-size-weighted correlation described above.

### Perturbation and TF prediction

#### Perturbation data generation

In silico perturbation was performed on the hematopoietic dataset across the 2,000 highly variable genes retained during preprocessing. Cells without valid lineage barcodes were removed, genes detected in fewer than ten cells were filtered out, and the count matrix was library-size normalized to 10,000 counts per cell, log-transformed, scaled and reduced to 50 principal components. The resulting PCA representation was used as the cell-state space for perturbation analysis.

For each highly variable gene, a gene-specific perturbation vector was constructed in PCA space using the corresponding PCA loading vector. The loading vector was normalized to unit length and scaled by the standard deviation of each principal component. The perturbation was applied along the positive PCA-loading direction, with the displacement defined as Δz = k(σPC ⊙ v), where v is the unit-normalized gene-loading vector, σPC is the standard deviation of each principal component, and k controls the perturbation magnitude. Because this operation shifts cells in PCA space rather than directly modifying a single gene, the perturbation should be interpreted as a gene-associated multivariate transcriptional displacement that may covary with other genes contributing to the same principal components.

A fixed perturbation magnitude was then applied to cells at the initial time point, corresponding to the early progenitor population. All other cells were left unchanged. This generated one perturbed input dataset for each screened gene, together with an unperturbed control dataset.

#### Perturbation screen

For each gene-specific perturbed dataset, cells at the initial time point were used as input particles and propagated forward using the trained LOTUS dynamics. An unperturbed control dataset was simulated in parallel using the same procedure. When simulated terminal states did not carry cell-type annotations, they were annotated using the classifier trained from observed single-cell states and cell-type labels.

For each target fate, downstream analysis was restricted to the initial cells or clones with measurable descendants in the relevant fate set. In the hematopoietic benchmark, for example, the analysis was restricted to initially undifferentiated cells whose lineage barcodes were observed among neutrophil or monocyte descendants at later time points. For each screened gene, the fraction of simulated terminal cells assigned to the target fate was calculated within the retained cell set. Perturbation effects were quantified as the difference in target-fate fraction between the perturbed and control simulations.

The perturbation screen was repeated ten times with different random seeds. In each repeat, the classifier was retrained, and both control and perturbed simulations were performed for every screened gene. For each gene and target fate, the mean control and perturbed fate fractions across repeats were used to calculate a log2 fold change. Statistical significance was assessed using a two-sided Welch’s t-test comparing perturbed and control fate fractions across repeats. P values were adjusted across genes using the Benjamini– Hochberg procedure.

For visualization and ranking, genes were summarized separately for each target-fate screen. Volcano plots were generated using the log2 fold change in target-fate fraction and the negative log10-transformed P value. Genes with significant FDR-adjusted P values were highlighted, and top-ranked genes were selected according to both effect size and statistical significance.

#### Perturbation comparison

To compare perturbation screens between LOTUS and the baseline method, screen results from different methods were analyzed using the same ranking and annotation procedure. For each target fate, genes were first filtered to retain positive regulators, defined as genes with a positive log2 fold change in the target-fate fraction and an FDR-adjusted P value below 0.05. Genes were then ranked by FDR-adjusted P value, nominal P value, negative log10-transformed P value and log2 fold change. The top-ranked genes from each method were extracted separately for each fate-specific screen.

To compare method-specific and shared perturbation candidates, the top-ranked genes identified by LOTUS and the baseline method were compared for each target fate to define three gene sets: LOTUS-only genes, baseline-only genes and genes shared by both methods. Shared genes were defined as genes present in the top-ranked lists of both methods, whereas method-specific genes were present only in one method’s list. This comparison was performed independently for each target fate.

To assess the biological specificity of perturbation-gene prioritization, top-ranked genes were annotated using curated fate-associated marker and transcription-factor sets. For the LARRY neutrophil and monocyte screens, target genes were defined as the union of lineage-associated markers and transcription factors, and their recovery was quantified across increasing ranking depths. For analyses comparing method-specific and shared candidates, genes were further classified according to their association with the target or alternative fate programs. The complete curated gene sets used for perturbation-screen annotation are provided in Supplementary Note 4.

#### TF prediction

To examine transcription-factor dynamics along LOTUS-inferred fate trajectories, day 2 cells were propagated forward using the trained LOTUS model and sampled at evenly spaced intermediate time points between the first day and the last day. The simulated trajectories were stored in the 50-dimensional PCA space together with their original lineage barcodes. For visualization of transcription-factor programs, simulated PCA coordinates were projected back to gene space using the PCA loading matrix.

### Details of other analysis

#### Lineage-tree construction

To summarize LOTUS-inferred differentiation dynamics at the population level, we constructed a lineage-tree representation by integrating cell-state geometry, inferred velocity, growth rate and potential. Cells without valid lineage barcodes were removed before analysis. Cell states were represented in the 50-dimensional PCA space used for model training. LOTUS-inferred potential, velocity and growth values were assigned to each cell for subsequent tree construction and visualization.

Cells were first grouped into local microclusters using a k-nearest-neighbor graph in PCA space followed by Leiden clustering. For each microcluster, we calculated the centroid in PCA space, the median potential-derived developmental stage, the majority cell-type label, the mean velocity magnitude and the mean inferred growth rate. The developmental stage was obtained by transforming the LOTUS-inferred potential into a normalized ordering, with early undifferentiated states placed near the root and more differentiated states placed progressively farther along the tree.

A directed backbone tree was then constructed between microclusters. Candidate edges were derived from the connectivity graph between neighboring microclusters. For each pair of connected microclusters, an initial structural score was computed from graph connectivity, cluster size and centroid distance. Edges were oriented from earlier to later developmental stages. When velocity information was used, the structural score was further weighted by the agreement between the mean source-cell velocity direction and the centroid displacement from the parent candidate to the child candidate. For each non-root microcluster, the parent was selected as the earlier neighboring microcluster with the highest velocity-weighted edge score. The root was selected from microclusters enriched for undifferentiated cells, prioritizing high undifferentiated-cell fraction, early developmental stage and large cluster size.

Tree coordinates were assigned after backbone construction. The vertical coordinate of each node was determined by its potential-derived developmental stage and constrained to increase monotonically from parent to child. Leaf nodes were ordered by post-order traversal, and horizontal coordinates were assigned so that internal nodes were positioned at the mean coordinate of their descendant leaves. Edges were rendered as smooth Bezier curves connecting parent and child nodes. Individual cells were assigned to the closest lineage path according to the distance between their microcluster centroid and the centroids along each root-to-leaf path, and were then positioned along the corresponding path according to their developmental stage.

Branch width was used to represent inferred local expansion. For each edge, a branch-strength score was calculated from the descendant subtree of the child node by summing cluster cell numbers weighted by the exponentiated normalized growth rate. Branch widths were scaled according to this score, whereas edge topology was determined by cell-state geometry and velocity alignment. Cell points were colored by annotated cell type, and terminal branches were labeled according to the majority cell type of the corresponding leaf microcluster.

#### Meta-clone construction

Meta-clones were defined by clustering clones with similar LOTUS-predicted trajectories from the initial time point. Clone similarity was quantified from normalized trajectory-occupancy profiles using Hellinger distance, followed by agglomerative clustering; the number of meta-clones was selected by silhouette score, yielding *k* = 10.

#### Multi-fate classification

To classify clone-level fate potential, day 2 cells were propagated forward by repeated stochastic simulation using the trained LOTUS model. Simulated states at later time points were annotated with the trained cell-type classifier, and the frequency of each predicted cell type across repeated simulations was used to estimate cell-level fate probabilities. These probabilities were then averaged across cells sharing the same lineage barcode to obtain clone-level fate probabilities.

For each clone, fate potential was determined from the non-undifferentiated cell types detected at later simulated time points. The probabilities from the intermediate and terminal time points were combined, and undifferentiated states were excluded. The number of remaining cell types with non-zero probability was defined as the clone’s fate number. Clones with one, two or more than two detected fates were classified as single-fate, double-fate or multi-fate clones, respectively; clones without any non-undifferentiated fate were classified as undifferentiated-only.

To quantify the agreement between predicted and observed clone-level fate-potential patterns, each clone was represented by a fate signature defined from the non-undifferentiated fates detected across the two later time points. Fate proportions at the two time points were averaged, and any fate with a mean proportion greater than 10⁻⁸ was included in the clone signature; signatures represented by fewer than five clones were grouped into an “other” category. We then compared the distributions of predicted and observed fate signatures using total variation (TV) distance, defined as TV(P,Q) = 1/2 Σ|P_i_ − Q_i_|, and Jensen–Shannon divergence (JSD), defined as JSD(P,Q) = 1/2 KL(P||M) + 1/2 KL(Q||M), where M = (P + Q)/2. JSD was normalized by ln(2) to obtain values between 0 and 1. TV-derived similarity and normalized JSD-derived similarity were defined as 1 − TV and 1 − JSD/ln(2), respectively, such that larger values indicate greater agreement between predicted and observed fate-signature distributions.

#### SCENIC regulon analysis

Regulon activity was analyzed using SCENIC. Gene regulatory networks were inferred using all barcoded cells, and cell-level regulon activity was quantified with AUCell. The AUCell matrix was then combined with clone-level fate-group annotations by lineage barcode. For the hematopoietic analysis, regulon AUC scores were compared among Neu-only, Mo-only and Neu/Mo double-fate cells. Differential regulon activity was assessed using the Kruskal–Wallis test, followed by Benjamini–Hochberg correction across regulons.

#### MultiVI-based ATAC imputation

Missing chromatin accessibility profiles for RNA-only query cells were imputed using MultiVI in scvi-tools version 1.3.3. Paired RNA–ATAC multiome cells were used as the reference dataset, and RNA-only query cells were added as expression-only observations. Query RNA features were aligned to the reference gene space, with missing genes filled with zeros. ATAC counts in the reference dataset were binarized before model training. The paired reference and RNA-only query data were combined using the MultiVI multiome data organization workflow. MultiVI was trained for up to 500 epochs with early stopping and a batch size of 128. After training, normalized chromatin accessibility probabilities were estimated for the RNA-only query cells using the trained model. The predicted accessibility matrix was exported as an AnnData object using the reference peak set.

#### Scribe temporal dependency analysis

Temporal gene-dependency networks were inferred using Scribe, which estimates whether past expression of a candidate regulator improves prediction of future target expression after conditioning on the target’s own history. LOTUS-reconstructed particles from the same clone and time point were collapsed into a single mass-weighted average expression state for Scribe input. For state-specific analyses, this aggregation was performed separately within each predicted TR1 state to generate clone-level temporal profiles. RDI was computed across multiple delays, and cRDI was used to reduce indirect dependencies by conditioning on additional upstream regulators. Separate effector, memory and activated TR1 networks were inferred using curated TF–target panels, and TF-source cRDI edges were interpreted as candidate state-specific temporal regulatory dependencies rather than direct binding interactions.

## Author contributions

J.L. designed and implemented the algorithm, performed the data analyses, prepared the figures and wrote the manuscript. H.H. performed the CD4⁺ T cell analyses, developed the website and wrote the manuscript.

Z.Z. contributed to the mathematical framework and algorithm design and wrote the theoretical sections. Yongjie L. provided advice on data preprocessing and downstream analyses. P.Z. and Yihan L. designed and supervised the project and reviewed and edited the manuscript.

## Acknowledgements

The authors acknowledge support from the National Natural Science Foundation of China under grants 92570207, 12288101, 8206100646, T2321001, 32550512, and T2325002, the Fundamental and Interdisciplinary Disciplines Breakthrough Plan of the Ministry of Education of China under grant JYB2025XDXM502, and the Fundamental Research Funds for the Central Universities

## Code and data availability

LOTUS is implemented in Python and is available at https://github.com/LinOT-Team/LOTUS (will be released alongside the paper). The raw data for the hematopoiesis dataset can be accessed at the Gene Expression Omnibus database with accession number GSE140802, the reprogramming dataset with accession number GSE99915. The raw data for the CD4+ T cell dataset are publicly available from the National Center for Biotechnology Information (BioProject PRJNA1129481).

## Supplementary Figures

**Supplementary Fig. 1.**
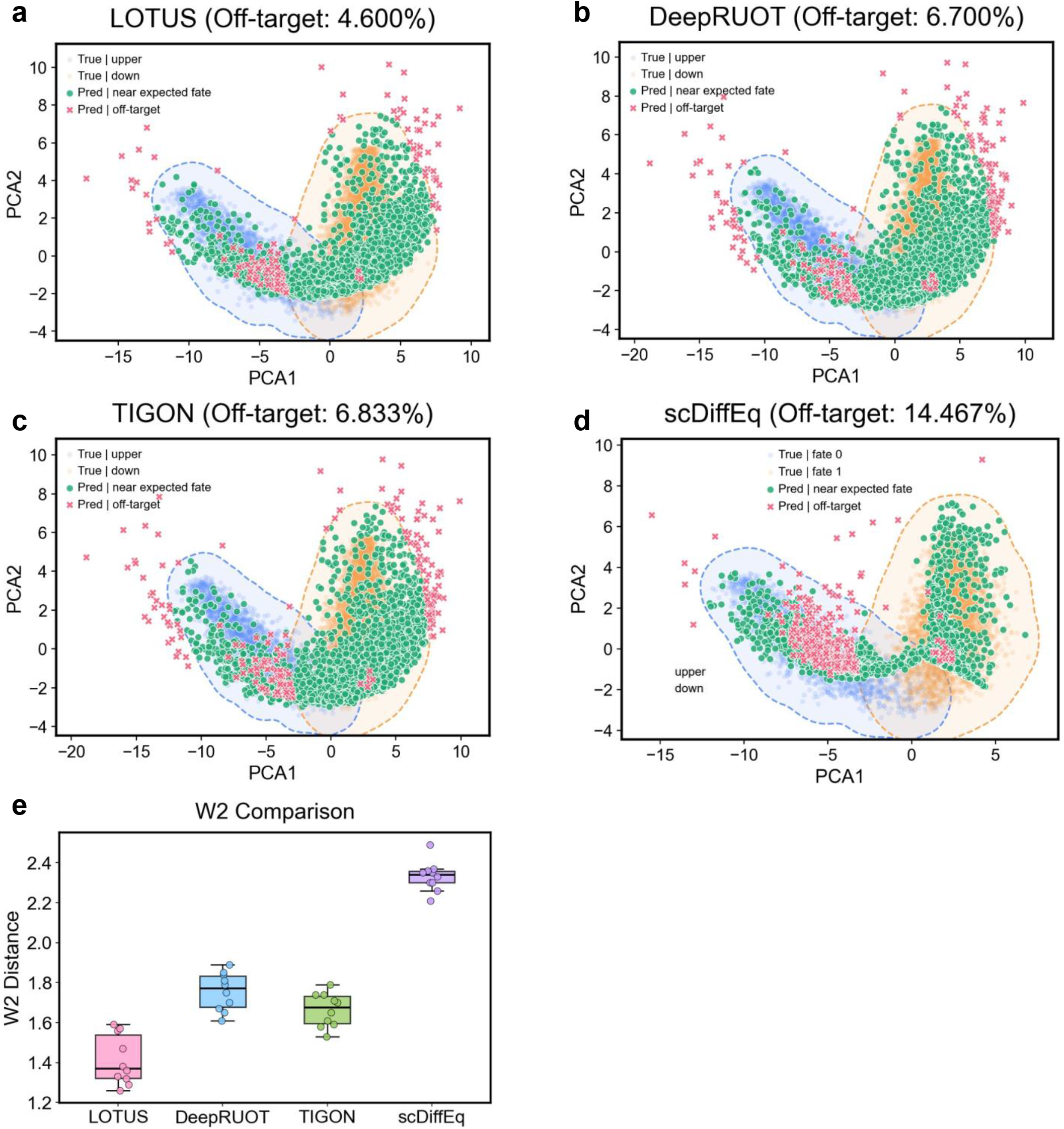
Benchmark of lineage-consistent trajectory reconstruction in simulated dataset 2. **a–d,** PCA visualizations of reconstructed trajectories for LOTUS (a), DeepRUOT (b), TIGON (c) and scDiffEq (d) in simulation dataset 2. True cells from the two barcode-associated fates are shown in blue and orange, correctly routed predicted cells are shown in green, and off-target predictions are shown in red. Dashed contours indicate the expected fate regions. **e,** Wasserstein-2 distance comparison across methods. Box plots show the final-time-point distance between predicted and observed distributions across ten simulations. Lower values indicate better agreement.

**Supplementary Fig. 2.**
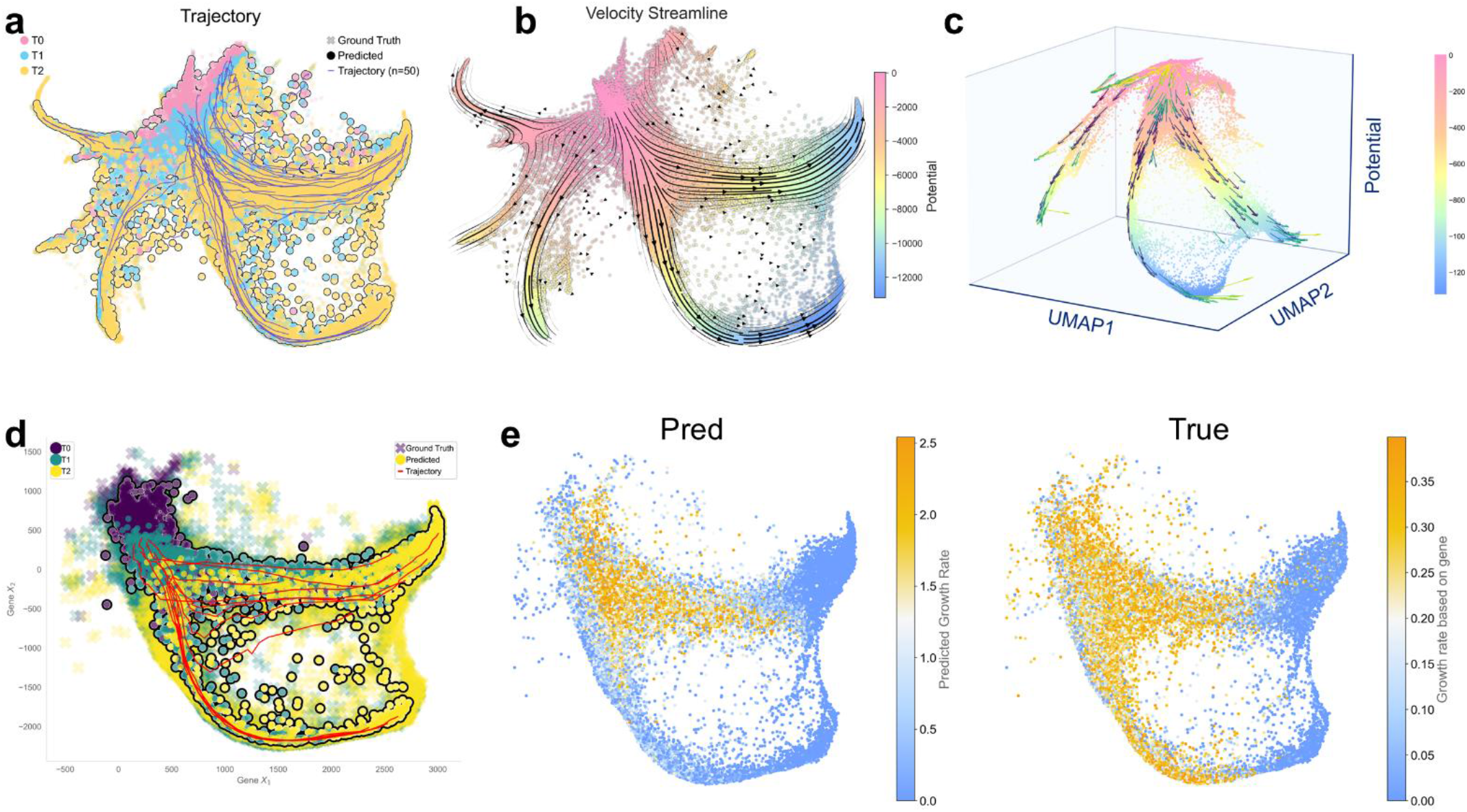
LOTUS-inferred trajectories, velocity field, differentiation potential and growth in the LARRY hematopoietic dataset. **a,** LOTUS-inferred stochastic trajectories projected onto the SPRING. Ground-truth cells from different time points are shown together with predicted cell states and sampled trajectories. **b,** Velocity streamlines inferred by LOTUS on the SPRING. Cells are colored by inferred differentiation potential. **c,** Three-dimensional visualization of the LOTUS-inferred potential landscape, with SPRING coordinates shown on the horizontal axes and inferred potential shown on the vertical axis. Streamlines indicate the direction of learned developmental flow. **d,** Representative LOTUS-generated trajectories in gene-expression space. Ground-truth cells from different time points are shown together with predicted cells and sampled trajectories. **e,** Comparison between LOTUS-predicted and reference growth-rate patterns on the SPRING. Colors indicate predicted growth rate or reference growth rate, showing the spatial organization of growth-associated expansion across the differentiation landscape.

**Supplementary Fig. 3.**
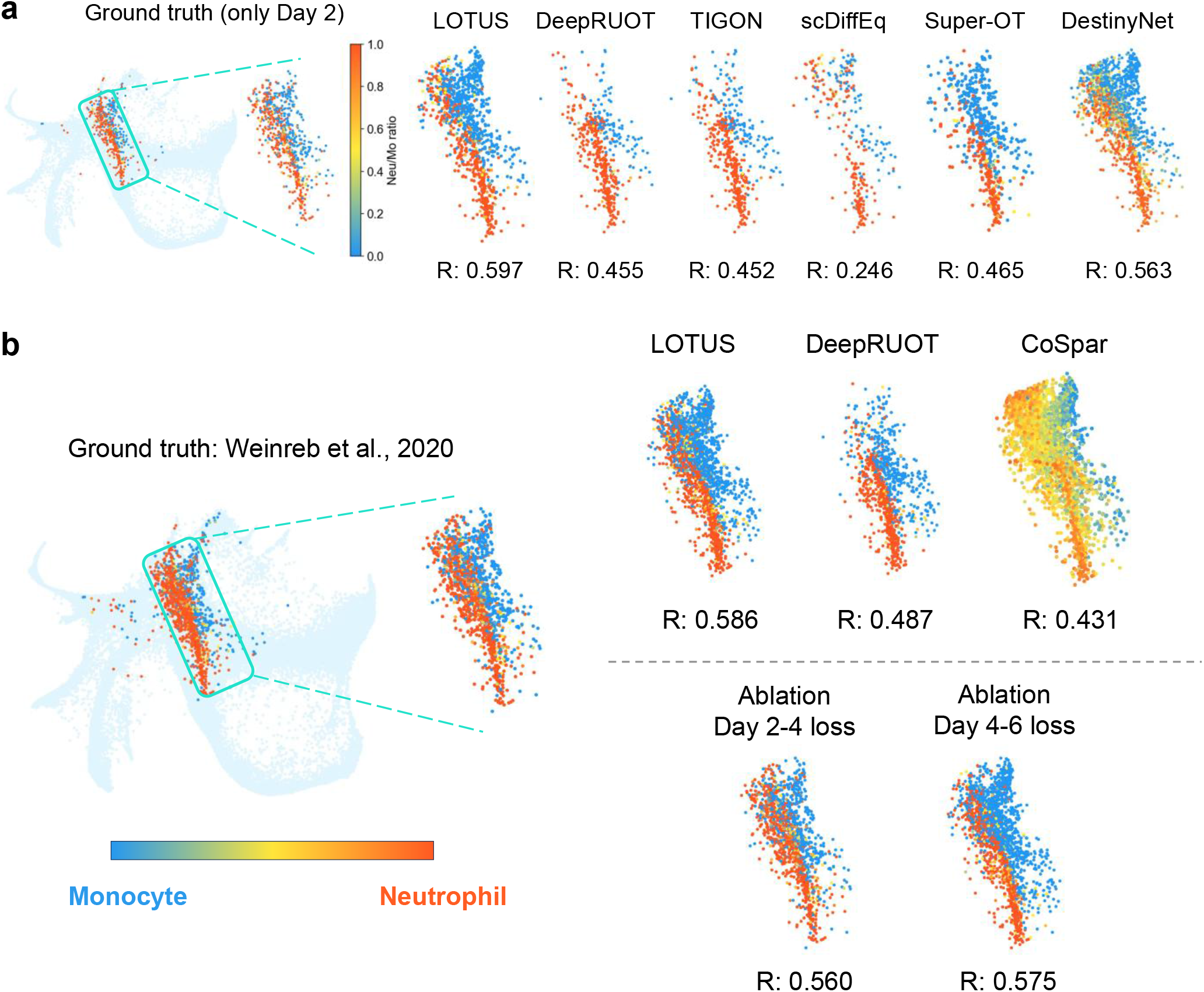
Local neutrophil–monocyte fate-boundary prediction in the LARRY dataset. **a,** Fate-boundary prediction between neutrophil and monocyte fates. Ground-truth neutrophil-to-monocyte fate ratios for Day 2 cells and predictions by LOTUS, DeepRUOT, TIGON, scDiffEq, Super-OT and DestinyNet are shown for the highlighted region. Colors indicate the neutrophil-to-monocyte fate ratio, and Pearson correlation coefficients *R* quantify agreement with the ground truth. **b,** Reference and model-predicted neutrophil–monocyte fate ratios in a boundary-associated region of the LARRY hematopoietic differentiation manifold. The left panel shows the ground-truth fate ratio derived from Weinreb et al. (2020), with the boxed region enlarged for quantitative comparison. Right panels show predicted fate-ratio maps from LOTUS, DeepRUOT and CoSpar, as well as LOTUS ablations with the Day 2–Day 4 or Day 4–Day 6 clone loss removed. Colors indicate the neutrophil-to-monocyte fate ratio, and Pearson’s r quantifies agreement with the ground-truth fate boundary.

**Supplementary Fig. 4.**
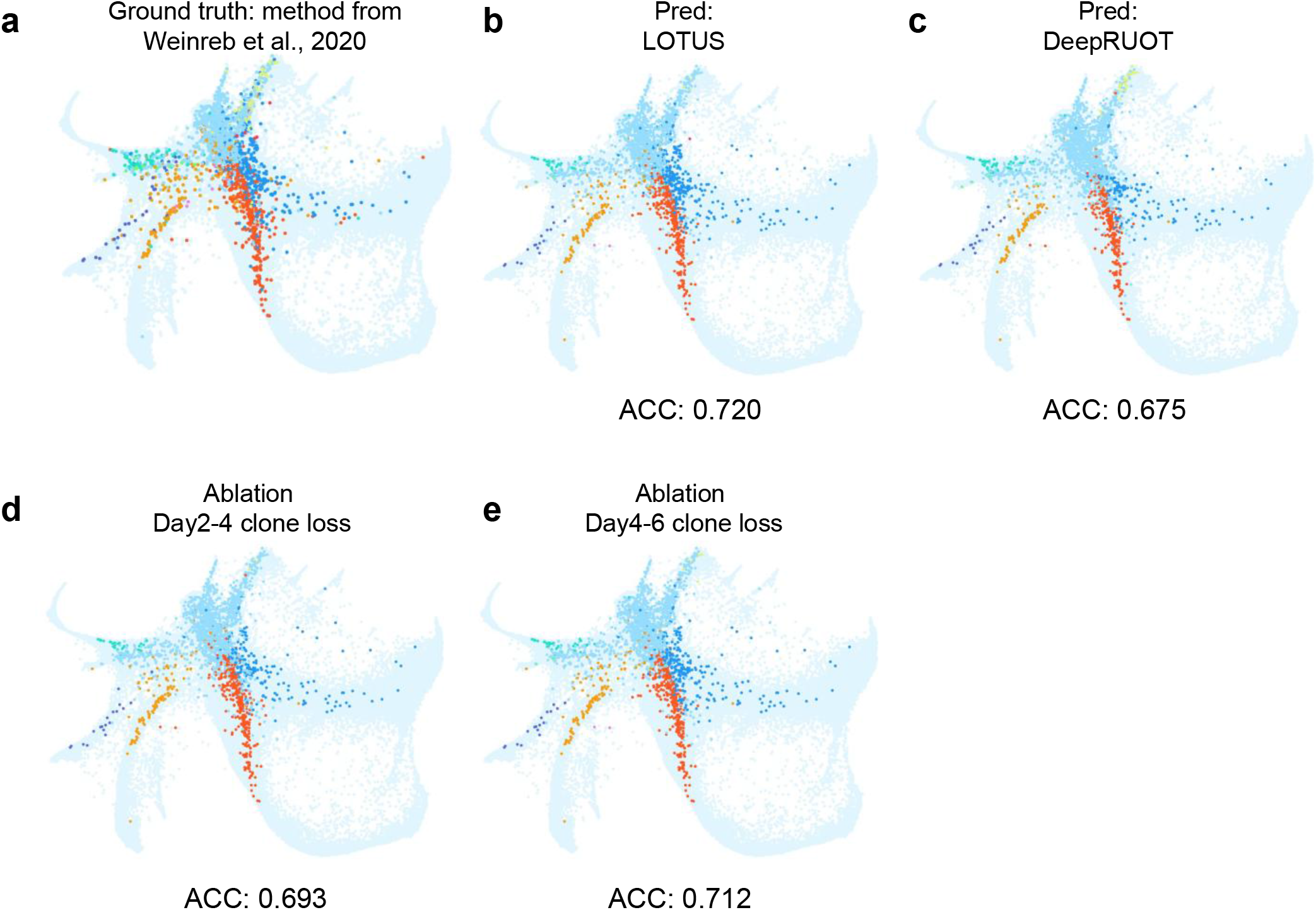
Benchmark of fate-label prediction and temporal lineage-supervision ablation in the LARRY dataset. **a,** Reference fate labels derived using the method from Weinreb et al. (2020). **b, c,** Predicted fate labels from LOTUS (**b**) and DeepRUOT (**c**) projected onto the SPRING. Accuracy values indicate agreement with the reference fate labels. **d, e,** Fate-label predictions after removing the day 2–day 4 clone loss (**d**) or the Day 4–Day 6 clone loss (**e**). Accuracy values quantify the effect of removing lineage supervision at each temporal transition.

**Supplementary Fig. 5.**
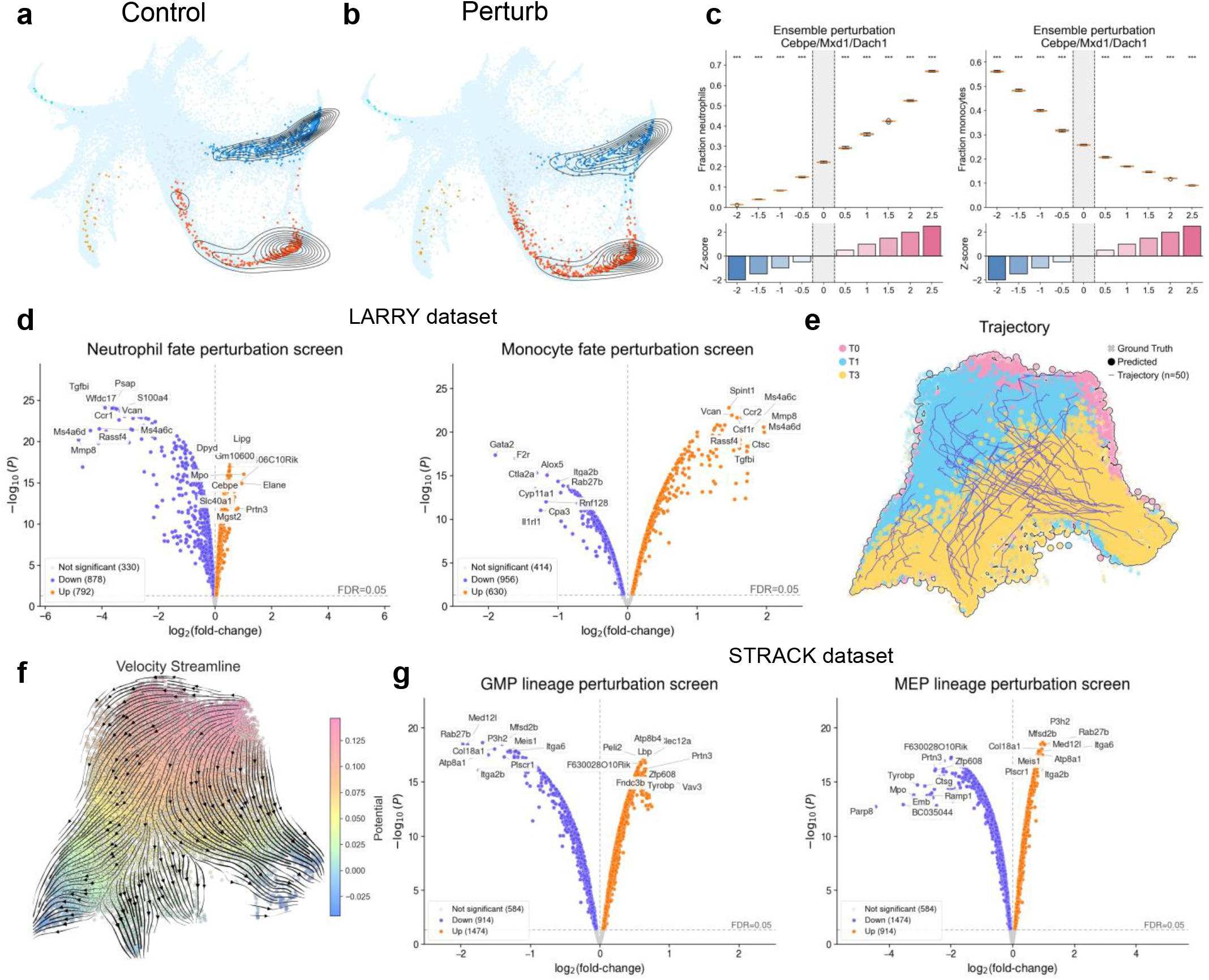
Additional *in silico* perturbation analyses and pathway enrichment in hematopoietic differentiation. **a, b,** LOTUS-inferred lineage-tree visualization on SPRING under control (**a**) and perturbation (**b**) conditions. The perturbation alters the predicted allocation of cells along neutrophil- and monocyte-associated branches. **c,** Dose-response analysis of ensemble perturbations involving *Cebpe*, *Mxd1* and *Dach1*. Line plots show changes in neutrophil and monocyte fractions across perturbation strengths, and lower bar plots show the corresponding perturbation z-scores. **d,** Volcano plots of neutrophil- and monocyte-associated perturbation screens in the LARRY dataset obtained through DeepRUOT. Significant genes with positive and negative effects on fate probability are highlighted, and top regulators are annotated. **e,** Comparison of predicted and ground-truth cell trajectories in the STRACK dataset. Colored trajectories show LOTUS-predicted lineage dynamics across the observed time points, while ground-truth endpoint states are overlaid for reference. **f,** Velocity streamline visualization of the inferred hematopoietic dynamics in the STRACK dataset. Streamlines indicate the direction of cell-state transitions, with the background colored by the inferred potential landscape. **g,** Volcano plots of GMP- and MEP-associated perturbation screens in the STRACK dataset obtained through DeepRUOT.

**Supplementary Fig. 6.**
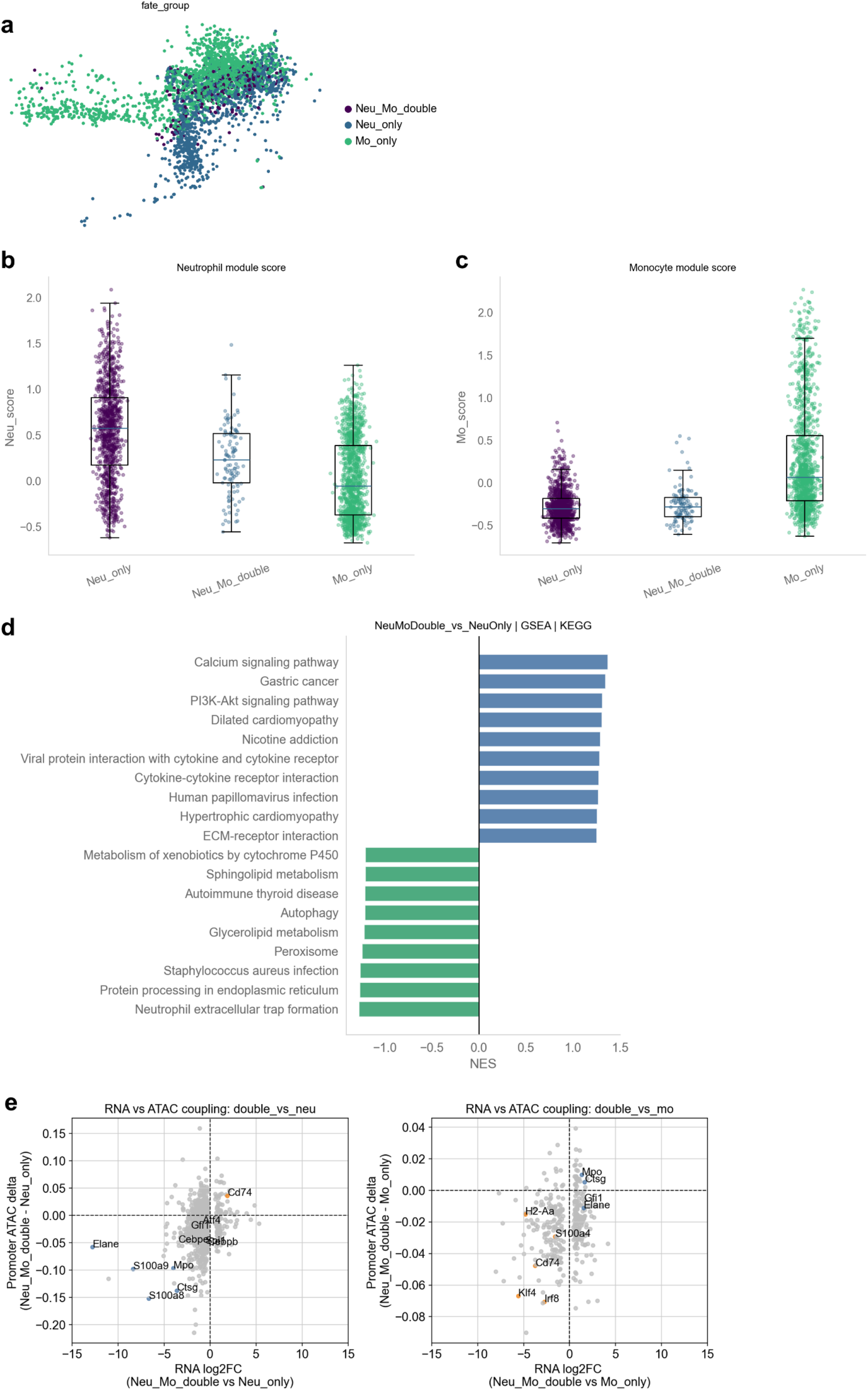
Molecular characterization of Neu–Mo double-fate cells. **a,** Distribution of Neu-only, Neu–Mo double-fate and Mo-only cells on the SPRING. **b, c,** Neutrophil module scores (**b**) and monocyte module scores (**c**) across Neu-only, Neu–Mo double-fate and Mo-only groups. Boxes indicate the interquartile range and center lines indicate the median. **d,** KEGG gene-set enrichment analysis comparing Neu–Mo double-fate cells with Neu-only cells. Bars show normalized enrichment scores for positively and negatively enriched pathways. **e,** Coupling between RNA expression changes and promoter ATAC accessibility differences in Neu–Mo double-fate cells relative to Neu-only or Mo-only cells. Each point represents a lineage-associated gene; labeled genes indicate representative concordant or discordant regulatory changes.

**Supplementary Fig. 7.**
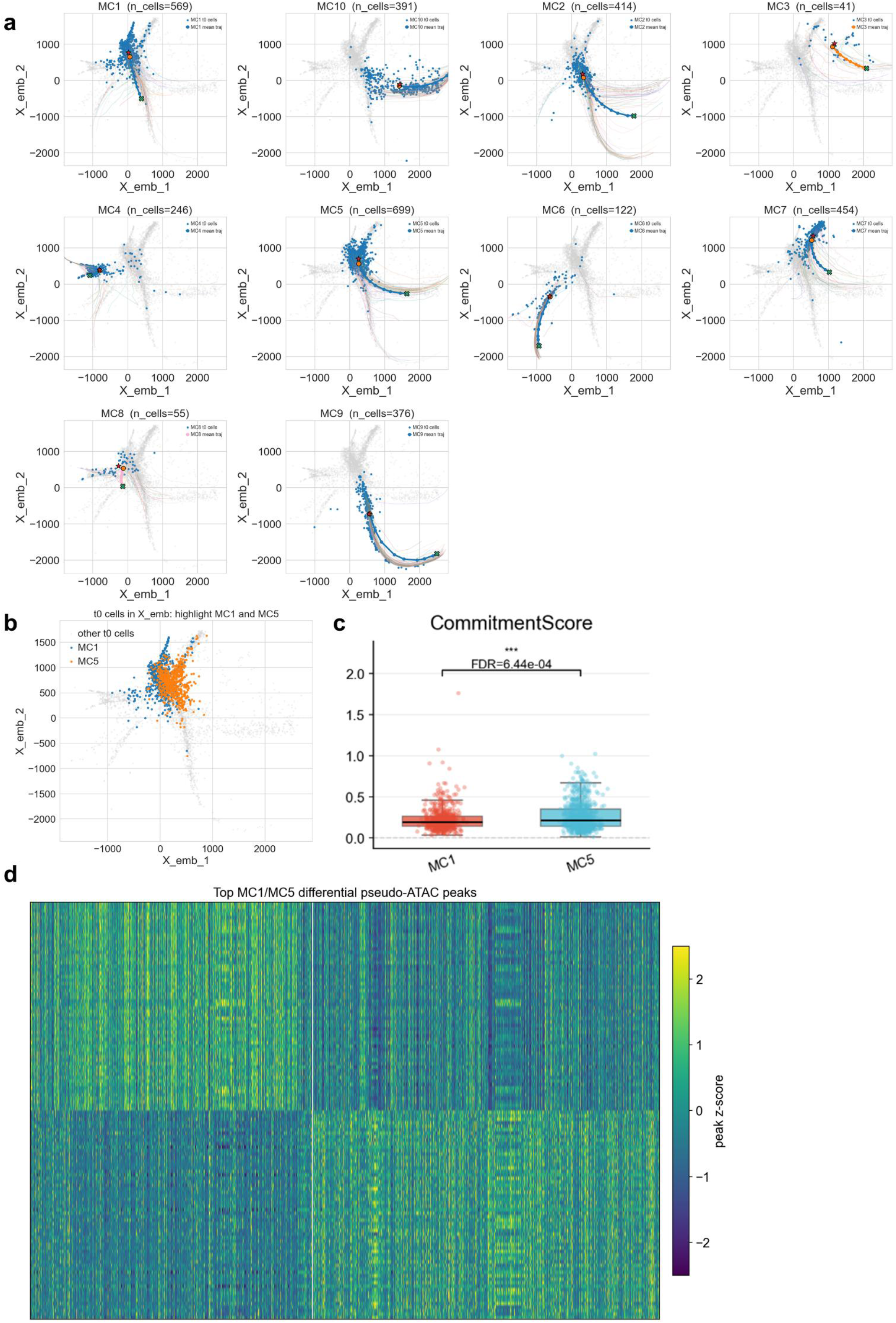
Meta-clone trajectories and regulatory differences. **a,** LOTUS-predicted trajectories for major meta-clone MC1–MC10 projected onto the initial cell embedding space. Blue points indicate cells belonging to each meta-clone, and colored trajectories show cluster-associated predicted differentiation routes. **b,** Distribution of MC1 and MC5 cells in SPRING. **c,** Boxplots showing the distribution of commitment scores in MC1 and MC5, with dots representing individual cells. Statistical significance was assessed by Mann–Whitney U test with FDR correction. The commitment score measures how strongly a cell is biased toward a single lineage program relative to alternative lineages. It was calculated from lineage-module accessibility scores, with higher values indicating stronger lineage commitment and lower values indicating broader multi-lineage priming. **d,** Heatmap of top MC1- and MC5-biased pseudo-ATAC peaks. Peaks were selected by the mean accessibility difference between MC1 and MC5, and values were z-scored across cells. Columns represent cells ordered by meta-clone identity, with the white line separating MC1 and MC5.

**Supplementary Fig. 8.**
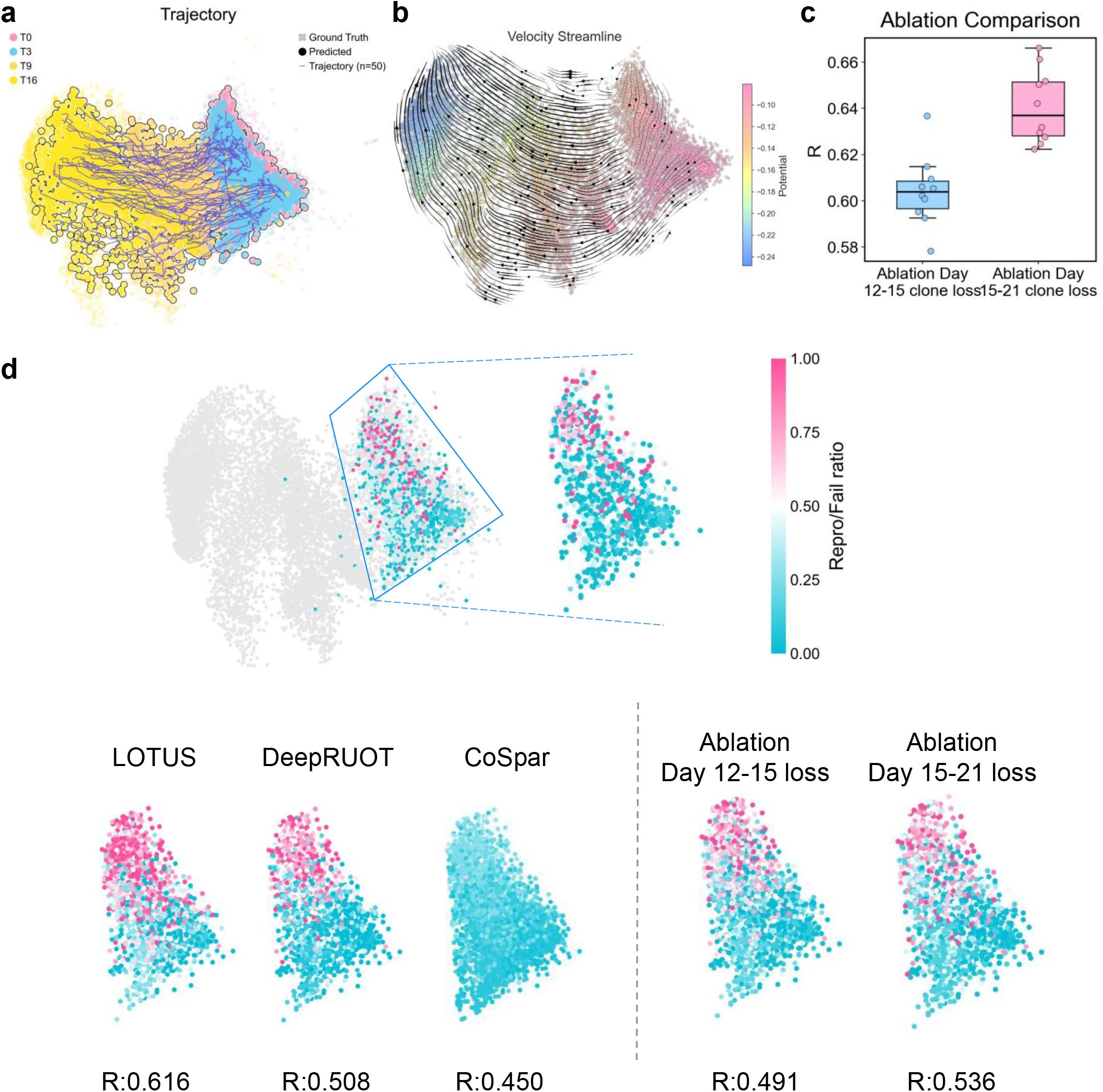
LOTUS reconstruction of reprogramming trajectories, velocity field and fate-boundary. **a,** LOTUS-inferred trajectories in the reprogramming dataset across sampled time points. Ground-truth cells, predicted cells and sampled trajectories are projected onto the UMAP. **b,** LOTUS-inferred velocity streamlines in the reprogramming dataset. Cells are colored by inferred potential, and streamlines indicate the direction of learned reprogramming dynamics. **c,** Pearson correlation between predicted and clone-derived reference fate ratios after ablating the day 12– 15 or day 15–21 clone loss (n = 10 independent runs). **d,** Top, reference reprogrammed-to-failed fate-ratio map in a boundary-associated region of the reprogramming UMAP. The boxed region is enlarged for quantitative comparison. Bottom, predicted fate-ratio maps from LOTUS, DeepRUOT and CoSpar, together with LOTUS ablations in which the Day 12→Day 15 or Day 15→Day 21 clone loss was removed. Colors indicate the reprogrammed-to-failed fate ratio, and Pearson’s r quantifies agreement with the reference fate-boundary structure.

**Supplementary Fig. 9.**
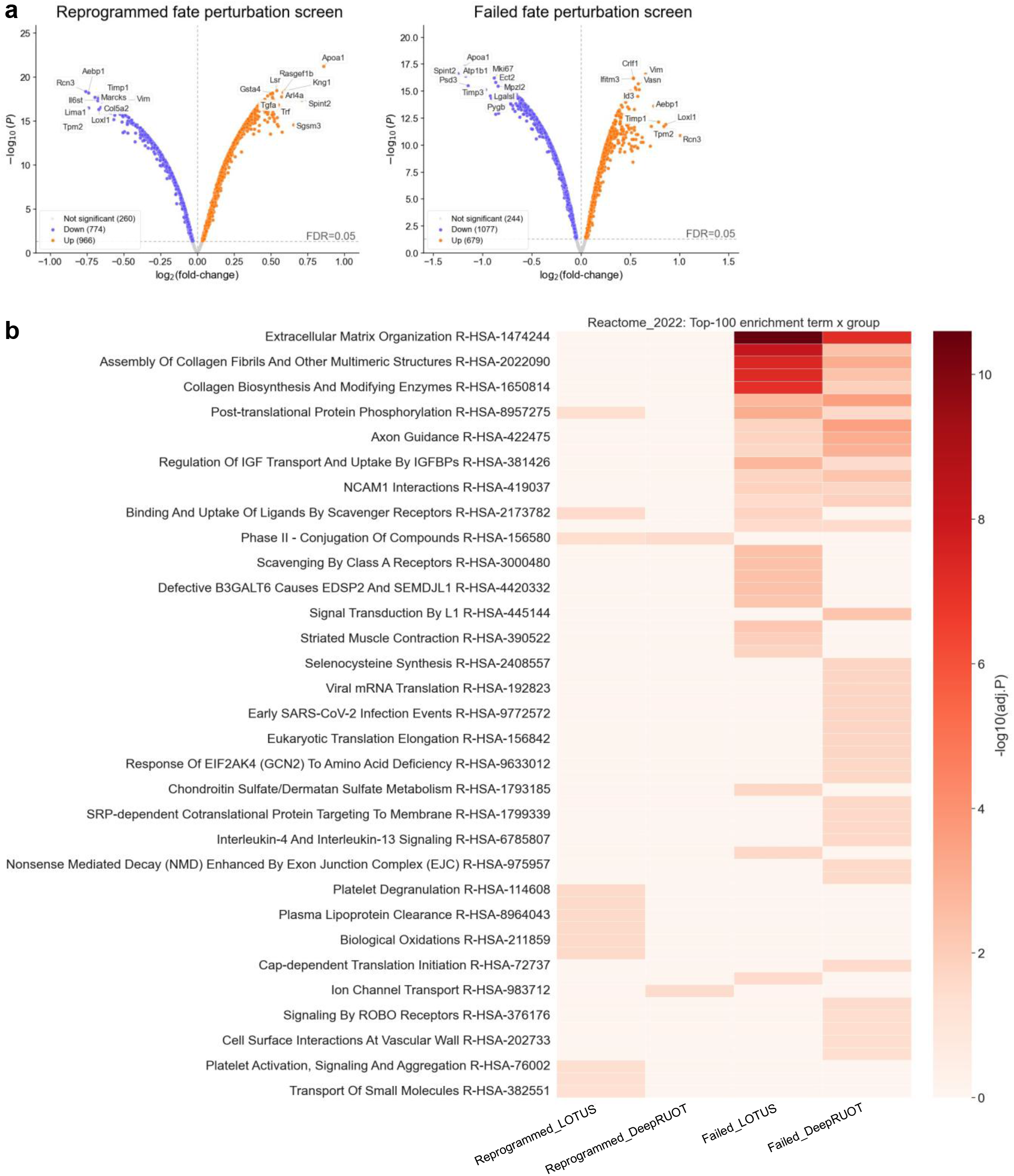
LOTUS perturbation-derived genes reveal fate-associated functional programs during reprogramming. **a,** Volcano plots showing predicted gene perturbation effects on reprogrammed and failed fate probabilities obtained through DeepRUOT. Significant genes with positive and negative effects on fate probability are highlighted, and top regulators are annotated. **b,** Reactome pathway enrichment heatmap of top-ranked perturbation genes from LOTUS and DeepRUOT in reprogrammed and failed fate screens. Color intensity represents enrichment strength, measured by −log10 adjusted *P* value.

**Supplementary Fig. 10.**
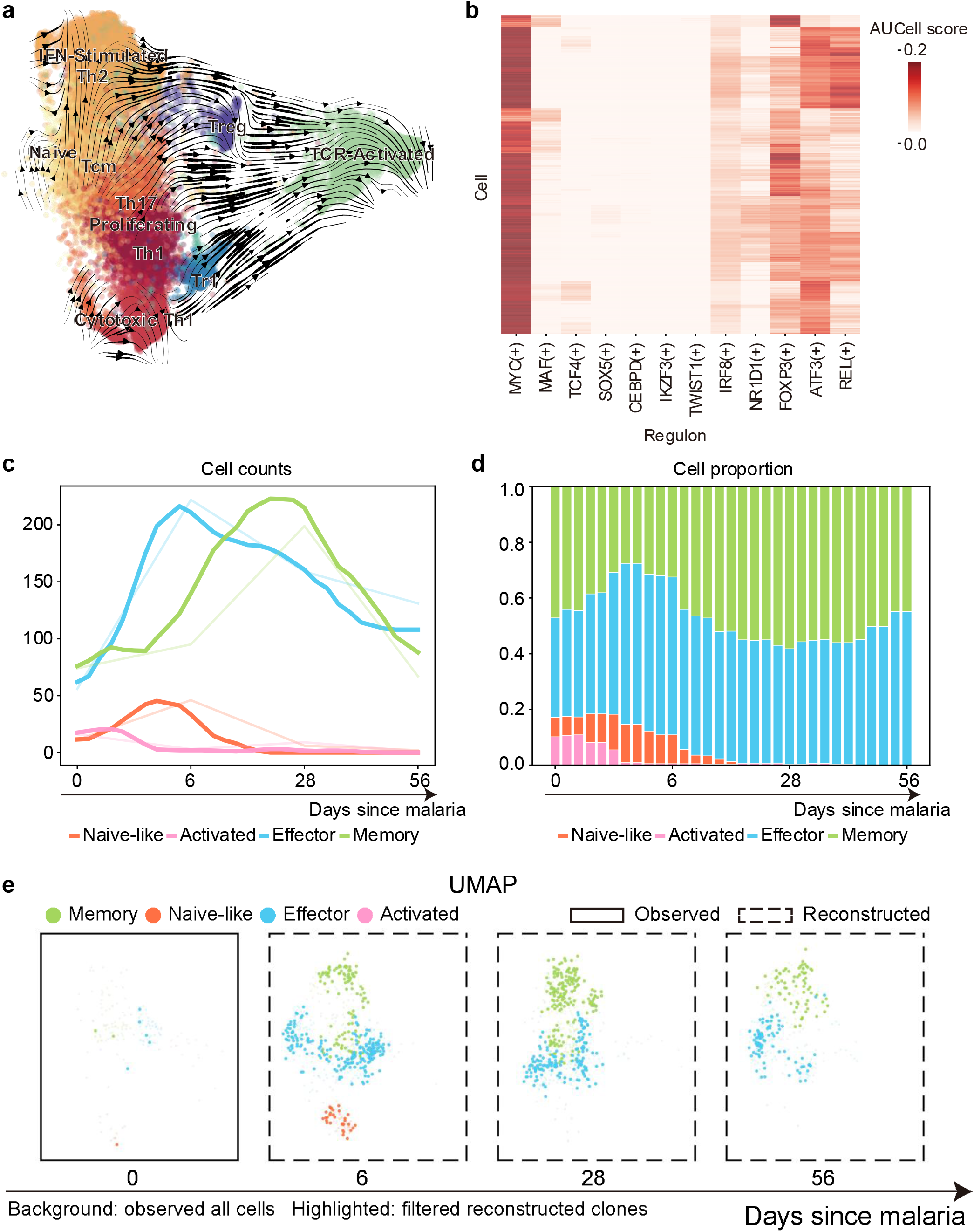
LOTUS resolves regulatory programs and continuous TR1 cell-state dynamics in TCR-activated CD4⁺ T cells during malaria infection. **a,** Velocity stream plot of LOTUS-reconstructed CD4+ T cells projected onto the reference UMAP embedding. Colors indicate CD4+ T-cell subsets, and black streamlines show the inferred direction of transcriptional state transition toward the TCR-activated population. **b,** pySCENIC AUCell regulon activity heatmap for LOTUS-reconstructed cells. Rows represent individual reconstructed cells and columns represent inferred regulons. Color intensity indicates AUCell score. **c**, Continuous temporal dynamics of LOTUS-reconstructed T_R_1 cell-state abundance across the course of malaria infection. Lines show the reconstructed abundance of naïve-like, activated, effector and memory T_R_1 cells over time after malaria infection. Pale lines indicate the corresponding trends in observed cell counts, whereas solid lines indicate smoothed LOTUS-reconstructed counts. **d**, Temporal changes in the proportions of LOTUS-reconstructed T_R_1 cell states across the infection course. Stacked bars show the relative proportions of naïve-like, activated, effector, and memory T_R_1 cells at each reconstructed time point. **e,** Reconstruction of longitudinal TR1 clonal dynamics in participant 3481 during malaria infection from clonotypes containing at least two cells. Background points indicate the observed distribution of all TR1 cells at each time point, whereas highlighted points represent LOTUS-reconstructed descendant cells from clonotypes containing at least two cells.

## References

1 Kester, L. & van Oudenaarden, A. Single-Cell Transcriptomics Meets Lineage Tracing. Cell Stem Cell 23, 166–179 (2018).

2 Wagner, D. E. & Klein, A. M. Lineage tracing meets single-cell omics: opportunities and challenges. Nature Reviews Genetics 21, 410–427 (2020).

3 Griffiths, J. A., Scialdone, A. & Marioni, J. C. Using single-cell genomics to understand developmental processes and cell fate decisions. Molecular Systems Biology 14, e8046 (2018).

4 Short, S., García-Tejera, R., Schumacher, L. J. & Coutu, D. L. Next generation lineage tracing and its applications to unravel development. Npj Systems Biology and Applications 11, 60 (2025).

5 Tang, F. C. et al. mRNA-Seq whole-transcriptome analysis of a single cell. Nature Methods 6, 377–386 (2009).

6 Trapnell, C. et al. The dynamics and regulators of cell fate decisions are revealed by pseudotemporal ordering of single cells. Nature Biotechnology 32, 381–386 (2014).

7 La Manno, G. et al. RNA velocity of single cells. Nature 560, 494–498 (2018).

8 Schiebinger, G. et al. Optimal-Transport Analysis of Single-Cell Gene Expression Identifies Developmental Trajectories in Reprogramming. Cell 176, 928–943.e22 (2019).

9 Tritschler, S. et al. Concepts and limitations for learning developmental trajectories from single cell genomics. Development 146, dev170506 (2019).

10 Weinreb, C., Wolock, S., Tusi, B. K., Socolovsky, M. & Klein, A. M. Fundamental limits on dynamic inference from single-cell snapshots. Proceedings of the National Academy of Sciences of the United States of America 115, E2467–E2476 (2018).

11 Weinreb, C., Rodriguez-Fraticelli, A., Camargo, F. D. & Klein, A. M. Lineage tracing on transcriptional landscapes links state to fate during differentiation. Science 367 (2020).

12 Mold, J. E. et al. Clonally heritable gene expression imparts a layer of diversity within cell types. Cell Systems 15, 149–165 (2024).

13 Carter, B. & Zhao, K. J. The epigenetic basis of cellular heterogeneity. Nature Reviews Genetics 22, 235–250 (2021).

14 Kim, M. & Costello, J. DNA methylation: an epigenetic mark of cellular memory. Experimental and Molecular Medicine 49, e322 (2017).

15 Mao, S. J. et al. Cell lineage tracing: Methods, applications, and challenges. Quantitative Biology 13, e70006 (2025).

16 Chen, C., Liao, Y. X. & Peng, G. D. Connecting past and present: single-cell lineage tracing. Protein & Cell 13, 790–807 (2022).

17 Zhang, L. et al. Lineage tracking reveals dynamic relationships of T cells in colorectal cancer. Nature 564, 268–272 (2018).

18 Hoehn, K. B. & Kleinstein, S. H. B cell phylogenetics in the single cell era. Trends in Immunology 45, 62–74 (2024).

19 Wang, S. W. MethylTree: exploring epimutations for accurate and non-invasive lineage tracing. Nature Methods 22, 463–464 (2025).

20 Ludwig, L. S. et al. Lineage Tracing in Humans Enabled by Mitochondrial Mutations and Single-Cell Genomics. Cell 176, 1325–1339 (2019).

21 Raj, B., Gagnon, J. A. & Schier, A. F. Large-scale reconstruction of cell lineages using single-cell readout of transcriptomes and CRISPR-Cas9 barcodes by scGESTALT. Nature Protocols 13, 2685–2713 (2018).

22 Biddy, B. A. et al. Single-cell mapping of lineage and identity in direct reprogramming. Nature 564, 219–224 (2018).

23 Pei, W. K. et al. Polylox barcoding reveals haematopoietic stem cell fates realized in vivo. Nature 548, 456–460 (2017).

24 Wang, S. W., Herriges, M. J., Hurley, K., Kotton, D. N. & Klein, A. M. CoSpar identifies early cell fate biases from single-cell transcriptomic and lineage information. Nature Biotechnology 40, 1066–1074 (2022).

25 Wang, K., He, X. L. & Hu, Z. Computational approaches for multimodal lineage tracing. Nature Reviews Genetics (2026).

26. Prasad, N., Yang, K. & Uhler, C. Optimal transport using GANs for lineage tracing. arXiv preprint arXiv:2007.12098 (2020).

27 Forrow, A. & Schiebinger, G. LineageOT is a unified framework for lineage tracing and trajectory inference. Nature Communications 12, 4940 (2021).

28 Lange, M. et al. Mapping lineage-traced cells across time points with moslin. Genome Biology 25, 277 (2024).

29 Liu, Z. et al. DestinyNet: A deep-learning framework for cell-fate analysis from lineage-tracing single-cell RNA sequencing data. Patterns 7, 101471 (2026).

30 Gao, M. Z. et al. CLADES: a hybrid NeuralODE-Gillespie approach for unveiling clonal cell fate and differentiation dynamics. Nature Communications 16, 8174 (2025).

31 Lin, Y., Chen, X., Wu, L., Zhou, Y. & Lin, Y. Widespread transcriptional memory shapes heritable states and functional heterogeneity in cancer and stem cells. Cell Systems, 101621 (2026).

32 Sha, Y. T., Qiu, Y. C., Zhou, P. J. & Nie, Q. Reconstructing growth and dynamic trajectories from single-cell transcriptomics data. Nature Machine Intelligence 6, 25–39 (2024).

33 Zhang, Z., Li, T. & Zhou, P. Learning stochastic dynamics from snapshots through regularized unbalanced optimal transport. International Conference on Learning Representations 2025, 19888–19919 (2025).

34 Zhang, Z., Wang, Z., Sun, Y., Li, T. & Zhou, P. Modeling cell dynamics and interactions with unbalanced mean field schrödinger bridge. Advances in Neural Information Processing Systems 38, 139518–139558 (2026).

35 Vinyard, M. E. et al. Learning cell dynamics with neural differential equations. Nature Machine Intelligence, 1–16 (2025).

36 Sashittal, P. et al. Inferring cell differentiation maps from lineage tracing data. Nature Methods 23, 532–541 (2026).

37 Yeo, G. H. T., Saksena, S. D. & Gifford, D. K. Generative modeling of single-cell time series with PRESCIENT enables prediction of cell trajectories with interventions. Nature Communications 12 (2021).

38 Bunne, C. et al. Learning single-cell perturbation responses using neural optimal transport. Nature Methods 20, 1759–1768 (2023).

39 Giladi, A. et al. Single-cell characterization of haematopoietic progenitors and their trajectories in homeostasis and perturbed haematopoiesis. Nature Cell Biology 20, 836–846 (2018).

40 Nakajima, H. & Ihle, J. N. Granulocyte colony-stimulating factor regulates myeloid differentiation through CCAAT/enhancer-binding protein ε. Blood 98, 897–905 (2001).

41 Korkmaz, B., Horwitz, M. S., Jenne, D. E. & Gauthier, F. Neutrophil Elastase, Proteinase 3, and Cathepsin G as Therapeutic Targets in Human Diseases. Pharmacological Reviews 62, 726–759 (2010).

42 Shi, C. & Pamer, E. G. Monocyte recruitment during infection and inflammation. Nature reviews immunology 11, 762–774 (2011).

43 Singh, I., Fernandez-Perez, D., Sanchez, P. S. & Rodriguez-Fraticelli, A. E. Pre-existing stem cell heterogeneity dictates clonal responses to the acquisition of leukemic driver mutations. Cell Stem Cell 32, 564–580 (2025).

44 Shivdasani, R. A., Fujiwara, Y., McDevitt, M. A. & Orkin, S. H. A lineage-selective knockout establishes the critical role of transcription factor GATA-1 in megakaryocyte growth and platelet development. Embo Journal 16, 3965–3973 (1997).

45 Canobbio, H., Balduini, C. & Torti, M. Signalling through the platelet glycoprotein Ib-V-IX complex. Cellular Signalling 16, 1329–1344 (2004).

46 Aibar, S. et al. SCENIC: single-cell regulatory network inference and clustering. Nature Methods 14, 1083–1086 (2017).

47 Ashuach, T. et al. MultiVI: deep generative model for the integration of multimodal data. Nature Methods 20, 1222–1231 (2023).

48 Zhang, Q. Y., Olofzon, R., Konturek-Ciesla, A., Yuan, O. Y. & Bryder, D. Ex vivo expansion potential of murine hematopoietic stem cells is a rare property only partially predicted by phenotype. Elife 12 (2024).

49 Shin, J. Y., Hu, W. H., Naramura, M. & Park, C. Y. High c-Kit expression identifies hematopoietic stem cells with impaired self-renewal and megakaryocytic bias. Journal of Experimental Medicine 211, 217–231 (2014).

50 Kamimoto, K. et al. Gene regulatory network reconfiguration in direct lineage reprogramming. Stem Cell Reports 18, 97–112 (2023).

51 Ross, J. et al. A Rare Human Syndrome Provides Genetic Evidence that WNT Signaling Is Required for Reprogramming of Fibroblasts to Induced Pluripotent Stem Cells. Cell Reports 9, 1770–1780 (2014).

52 Watanabe, K. et al. OVOL2 induces mesenchymal-to-epithelial transition in fibroblasts and enhances cell-state reprogramming towards epithelial lineages. Sci Rep. 9, 6490 (2019).

53 Xing, Q. R. et al. Diversification of reprogramming trajectories revealed by parallel single-cell transcriptome and chromatin accessibility sequencing. Science Advances 6, eaba1190 (2020).

54 Nideffer, J. et al. Clone tracking through repeated malaria identifies high-fidelity memory CD4 T cell responses. Science Immunology 10, eads2957 (2025).

55 Qiu, X. J. et al. Inferring Causal Gene Regulatory Networks from Coupled Single-Cell Expression Dynamics Using Scribe. Cell Systems 10, 265–274 (2020).

56 Karwacz, K. et al. Critical role of IRF1 and BATF in forming chromatin landscape during type 1 regulatory cell differentiation. Nature Immunology 18, 412–421 (2017).

57 Nideffer, J. & Jagannathan, P. Type I regulatory T cells in malaria: of mice and men. The Journal of Clinical Investigation 133 (2023).

58. Lipman, Y., Chen, R. T., Ben-Hamu, H., Nickel, M. & Le, M. Flow matching for generative modeling. arXiv preprint arXiv:2210.02747 (2022).

59. Tong, A., et al. Improving and generalizing flow-based generative models with minibatch optimal transport. arXiv preprint arXiv:2302.00482 (2023).

60 Tong, A. et al. Simulation-free Schrödinger bridges via score and flow matching. arXiv preprint arXiv:2307.03672 (2023).

61 Jindal, K. et al. Single-cell lineage capture across genomic modalities with CellTag-multi reveals fate-specific gene regulatory changes. Nature Biotechnology 42, 946–959 (2024).

62 Metzner, E., Southard, K. M. & Norman, T. M. Multiome Perturb-seq unlocks scalable discovery of integrated perturbation effects on the transcriptome and epigenome. bioRxiv : the preprint server for biology (2024).

63 Li, L., Bowling, S., Wang, S. W. & Camargo, F. A mouse model with high clonal barcode diversity for joint line-age, transcriptomic, and epigenomic profiling in single cells. Experimental Hematology 124, S44–S44 (2023).

64 Wolf, F. A., Angerer, P. & Theis, F. J. SCANPY: large-scale single-cell gene expression data analysis. Genome biology 19, 15 (2018).

